# ecTMB: a robust method to estimate and classify tumor mutational burden

**DOI:** 10.1101/602060

**Authors:** Lijing Yao, Yao Fu, Marghoob Mohiyuddin, Hugo YK Lam

## Abstract

Tumor Mutational Burden (TMB) is a measure of the abundance of somatic mutations in a tumor, which has been shown to be an emerging biomarker for both anti-PD-(L)1 treatment and prognosis. Nevertheless, multiple challenges still hinder the adoption of TMB for clinical decision-making. The key challenges are the inconsistency of TMB measurement among assays and a lack of meaningful threshold for TMB classification. We describe a powerful and flexible statistical framework for estimation and classification of TMB (ecTMB). ecTMB uses an explicit background mutation model to predict TMB robustly and consistently. In addition to the two known TMB subtypes, TMB-high and TMB-low, we discovered a novel TMB subtype, named TMB-extreme, which was significantly associated with patient survival outcome. This discovery enabled ecTMB to classify samples to biologically and clinically relevant subtypes defined by TMB.

## Introduction

Research has shown that cancers can be caused by an accumulation of genetic mutations in oncogenes or tumor suppressors^1^. These mutations are known as “driver” mutations and they are under positive selection. However, only a very small fraction of somatic mutations in a tumor sample are expected to be drivers. The remaining majority of somatic mutations are “passengers,” accumulated randomly with a background mutation rate (BMR) during cancer progression^2^. Moreover, it has been shown that the somatic mutation rates of cancer patients vary^3^. A patient with a high somatic mutation rate is referred to having a hypermutated phenotype. Environmental exposure^4^, DNA synthesis/repair dysfunction^3,5^, and regional mutation heterogeneity^3,6^ account for the mutational rate heterogeneity. For example, deleterious mutations in *POLE*, *POLD1*, and the MMR system defects may lead to a hypermutated phenotype^3,7^. Seven genes have been identified as essential components for MMR system, including *MLH1*, *MLH3*, *MSH2*, *MSH3*, *MSH6*, *PMS1*, *PMS2*^8^.

Recently, immunotherapies targeting immune checkpoint inhibitors, such as programmed cell death protein 1 (PD-1), along with its receptor (PD-L1), and cytotoxic T lymphocyte-associated antigen 4 (CTLA-4), have shown remarkable clinical benefits for various advanced cancers^9^. However, only a fraction of patients is responsive to the treatment, making it critical to identify predictive biomarkers to distinguish responsive patients. PD-L1 expression level and microsatellite instability-high (MSI-H) have been developed to be predictive biomarkers for anti-PD-L1 therapy^10^. Microsatellite instability (MSI) is a phenotype of an accumulation of deletions/insertions in repetitive DNA tracts, called microsatellites. Similar to hypermutation, evidences have indicated that MSI results from a deficient MMR system^11^.

Hypermutation was first associated with the response to CTLA-4 blockade therapy in 2014^12^ and PD-1 blockade therapy in 2015^13^. Since then, tumor mutational burden (TMB), which is a measure of the abundance of somatic mutations, has become a new promising biomarker for both prognosis^14^ and immunotherapy^15^. Nevertheless, multiple challenges still hinder the adoption of TMB for clinical decision making. The current well-accepted TMB measurement requires counting the non-synonymous somatic mutations in a paired tumor-normal sample using whole-exome sequencing (WES). However, clinical diagnostics based on sequencing technologies still rely heavily on targeted panel sequencing. Although studies have shown panel-based TMB measurements were highly correlated with WES-based TMB^15–17^, inconsistencies between these two measurements have been observed^16–19^. One reason for this inconsistency is that targeted panel sequencing might overestimate TMB due to its enrichment of driver mutations and mutation hot spots. In fact, WES-based TMB is more indicative of the overall BMR because of the lower incidence of driver mutations in whole exome. In order to avoid overestimating TMB, various filtering strategies have been applied. For example, Foundation Medicine used COSMIC to filter out driver mutations and add synonymous mutations to reach an agreement with WES-based TMB^16^. These arbitrary filters are dependent on frequently updated databases, worsening the inconsistency, reproducibility and robustness of the calculation. Another non-negligible challenge is the relatively arbitrary selection of the TMB-high cutoff such as 10 or 20 per Mb or top 10% or 20% quantile^20^. Although these thresholds were enough to illustrate the predictive value of TMB as a biomarker, an appropriate TMB cutoff derived from sophisticated studies or clinical trials would be better.

In order to improve the robustness of TMB measurement and TMB subtype classification, we proposed a novel method called ecTMB (estimation and classification of TMB) (Fig. 1). Because WES-based TMB is akin to the overall BMR, we built a statistical model using a Bayesian framework for TMB prediction. Furthermore, log transformation of the TMB revealed three TMB subtypes: TMB-low, TMB-high and a novel subtype – TMB-extreme. Based on this observation, we extended ecTMB with a Gaussian Mixture Model to classify samples by the aforementioned cancer subtypes. Our method was evaluated using WES data from The Cancer Genome Atlas (TCGA) as describe in method.

**Fig. 1.**
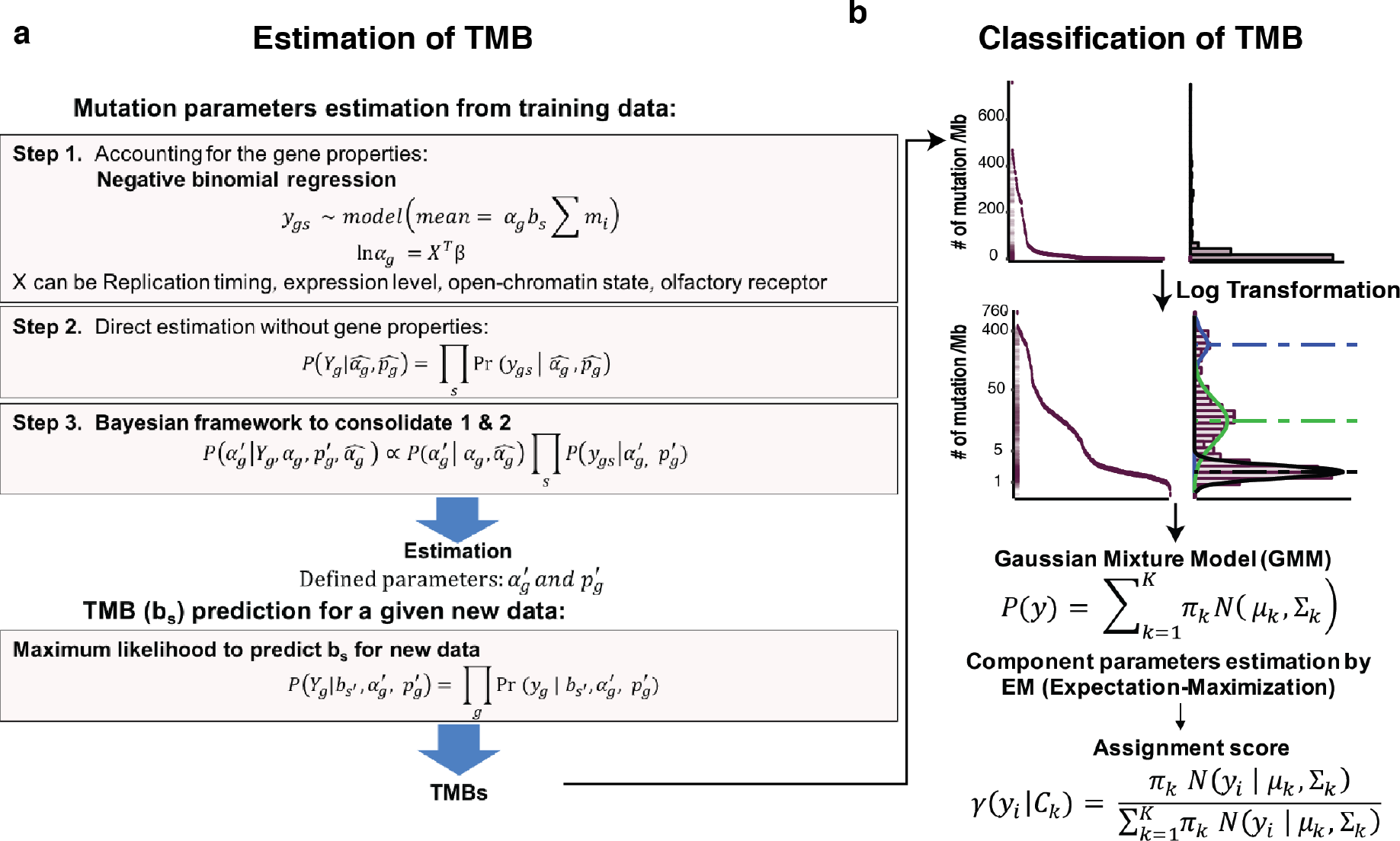
ecTMB workflow. (a) An explicit background mutation model for TMB prediction. (b) TMB subtype classification based on log-transformed TMB using GMM.

## Results

### Background mutation modeling

BMR modeling is one of the major challenges for driver mutation detection. Multiple methods have been developed to model BMR. MutSigCV^3^ applies genomic features to estimate BMR and DrGaP builds a Bayesian framework to take 11 mutation types into consideration for BMR estimation^21^. However, cancer mutational heterogeneity is far more complex. Here, we assumed the occurrence of silent mutations follow the BMR without selection pressures and the number of background somatic mutations follows a negative binomial distribution. In order to incorporate all known factors, e.g. tri-nucleotide context^5^, sample-specific mutational burden, gene expression level^3^ and replication timing^22^, the Generalized Linear Model (GLM) was used to estimate the general impact of these factors by pooling genes together (Fig. 1A). In order to evaluate our model, we divided samples corresponding to each cancer type into training and test sets with a 70% :30% split. Training sets ware used to estimate the model parameters, which then were used to predict the number of mutations for each gene of each sample based on the negative binomial distribution (see Methods). Because of the assumption that synonymous mutations were accumulated with a BMR, the comparison of predicted number of synonymous mutations against observed can be used to measure the performance of model. We found that the GLM model could not explain all variations in the observed number of synonymous mutations. For example, membrane-associated mucin (MUC16) and titin (TTN), which are two suspicious false-positive driver genes^3^, had much lower predicted number of synonymous mutations than actually observed in both training and test sets (Supplementary Fig. 1). Therefore, there might be unknown sequencing or biological factors influencing BMR as well.

To handle unknown factors, we separately modeled each gene as an independent negative binomial process as the second step. The final adjusted gene-specific BMRs were then generated through a Bayesian framework to consolidate the estimators from the previous two steps (Fig 1A) (details in Methods). Compared with prediction of synonymous mutations from GLM, the final model improved R-squared from 0.5 to ~0.9 in the training set and from 0.3 to ~ 0.6 in the test set and reduced the mean absolute error (MAE) and the root mean square error (RMSE). Meanwhile, mutation predictions for MUC16 and TTN become much closer to observed values (Supplementary Fig. 1). These results displayed an improved performance when our three-step approach was applied.

A driver gene was expected to possess a higher non-synonymous mutation frequency relative to its BMR due to the positive selection. Indeed, we discovered several well-known cancer-specific driver genes whose observed number of non-synonymous mutations are much higher than predicted background ones. Examples of such driver genes includes: TP53, KRAS, PIK3CA and SMAD4 in colorectal cancer^23^, TP53, ARID1A and PIK3CA in stomach cancer^24^, and PTEN, ARID1A, PIK3CA and TP53 in endometrial cancer^25^ (Supplementary Fig. 1). In summary, these results demonstrated that our method can accurately model background mutations, systematically reducing driver gene impacts.

### TMB prediction

In our model, there are three determinants for the BMR, namely: sequence composition, gene-specific BMR and sample-specific BMR. As with the training process described above, gene-specific BMRs can be estimated under the assumption that the sample-specific BMR of a sample could be calculated as the number of non-synonymous mutations per Mb, which is equivalent to TMB. With determined gene-specific BMRs from training set as described above, sample-specific BMR for a new sample could be estimated by Maximum Likelihood Estimation (MLE) through modeling each gene as an independent negative binomial process (Fig. 1A, detail in Method).

Using test sets, we firstly evaluated how good TMB predictions by ecTMB were when all mutations, non-synonymous as well as synonymous, from WES were used. ecTMB was compared against the standard TMB measurement (WES-based TMB), calculated by the number of non-synonymous mutations divided by sequenced genomic region size. Pearson correlation coefficient (R) is widely used to assess the agreement of TMB measurements among assays. However, a high correlation does not mean two methods agree because R measures the strength of a relation between two variables but not the exact agreement between them^26^. In order to comprehensively assess the agreement, we not only used correlation coefficient, but also measured MAE, RMSE and performed Bland-Altman analyses. We found that the TMB predictions by ecTMB were highly concordant with standard TMB calculations for both correlation (correlation coefficient > 0.998) and absolute error (MAE < 1.833 in linear scale and MAE < 0.063 in log scale) metrics.

ecTMB can use synonymous mutations for TMB prediction since synonymous mutations follow the background mutation accumulation. Meanwhile, it is also able to incorporate non-synonymous mutations, most of which follow the BMR as well. We further assessed the impact of including non-synonymous mutations from different proportions of genes. Genes were ranked based on mutation frequency in training set for each cancer type and non-synonymous mutations from the least mutated genes (bottom 0%, 20%, 60%, 80%, 85%, 90%, 95% and 100%) were added to the prediction. In all, while predictions with only synonymous mutations already achieved a great concordance with WES-based standard TMB with R > 0.975 and almost 0 bias, the addition of non-synonymous mutations further improved the concordance, where R > 0.999 and 0 bias (Supplementary Fig. 2).

**Fig. 2.**
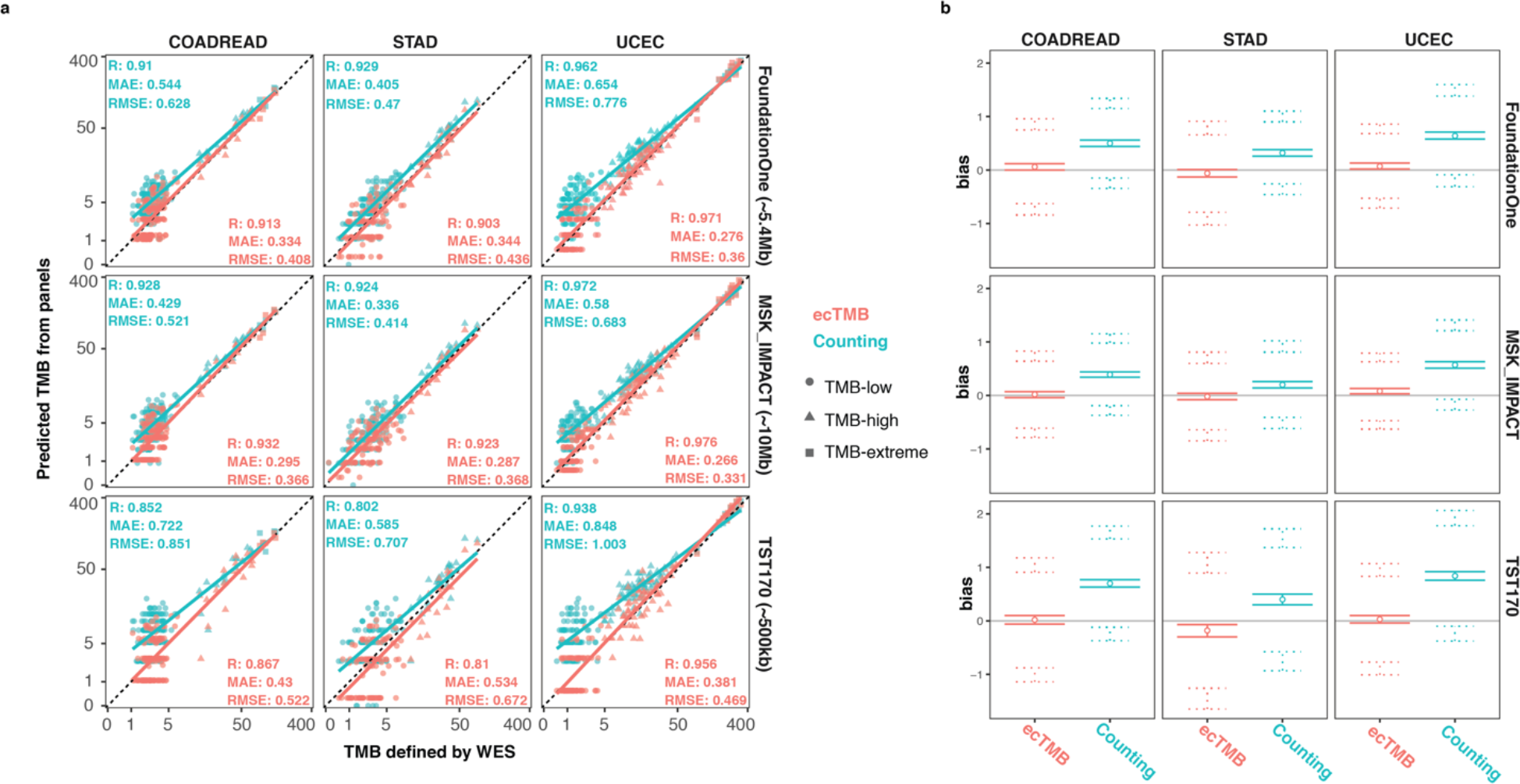
Evaluation of the panel-based TMB prediction performance. (a) Scatter plots show WES-based standard TMB plotted against predicted panel-based TMBs for each cancer type and panel combination. Two methods were used for panel-based TMB predictions, including counting method (in cyan) and ecTMB (in red). Their linear regression lines against WES-based TMB and performance measurements (correlation coefficient, MAE and RMSE) are plotted for each method in each scatter plot. (b) Bland Altman analysis results for counting method (cyan) and ecTMB (red) against WES-based TMB are shown. The middle circle indicates the bias (mean difference) and two solid lines around it are 95% confidence interval for the bias. The two dotted line on the top are 95% confidence intervals for the upper limit of 95% agreement and ones on the bottom are 95% confidence intervals for the lower limit of 95% agreement.

We further conducted *in silico* assessments of panel-based TMB prediction by the counting method and ecTMB on three cancer panels, including FoundationOne CDx, Integrated Mutation Profiling of Actionable Cancer Targets (MSK-IMPACT)^27^ and Illumina TruSight Tumor 170 (TST170). As with other studies^15–17^, we detected a high correlation between WES-based standard TMB and panel-based TMB through simply counting the number of non-synonymous mutations. But Bland-Altman analyses showed the significant biases (> 0) of counting method, indicating an over-estimation especially for low TMB samples (Fig. 2 and Supplementary Fig. 3). Samples with low TMB were more vulnerable to over-estimation since fewer background mutations led to a higher representation of cancer-associated mutations in counting. In contrast, ecTMB predictions, using synonymous and 95% of non-synonymous mutations, not only had comparable or improved correlation coefficients with WES-based TMB, but also reduced MSE, RMSE and biases. As an example, for the predictions of TST170 panel in endometrial cancer, ecTMB improved correlation coefficient from 0.938 to 0.956, reduced MAE from 0.848 to 0.381 and removed bias (from 0.84 with 95% confidence interval [0.76, 0.92] to 0.03 [−0.04, 0.1]), compared with counting prediction (Fig. 2). The reasons for using 95% of non-synonymous mutations were 1) fewer synonymous mutations detected within each panel led to less accurate predictions; 2) too many driver gene mutations resulted in prediction biases (Supplementary Fig. 3). In fact, the mean number of synonymous mutations in colorectal cancer were 4.83, 5.67, 3.55 for FoundationOne, MSK-IMPACT and TST170 panel, respectively.

### Three cancer subtypes revealed by log-transformed TMB

While exploring the distribution of TMB, we discovered that the distribution of log-transformed WES-based TMB, defined by either the number of all mutations or non-synonymous mutations per Mb, resembled a mixture of gaussians in colorectal, stomach and endometrial cancers (Fig. 3 and Supplementary Fig. 5). We extended our investigation of this observation to all cancer types in TCGA by grouping cancer types based on the similarity of log-transformed TMB distribution as described in Methods (Supplementary Fig. 6). However, we could not identify the same pattern in those groups, which might be due to very few hypermutated samples, such as groups 1 and 5, or due to environmental factors causing a continuous mutation spectrum, such as group 2 (Supplementary Fig. 7). Because of the lack of same pattern in those cancer types, we focused our analyses only on colorectal, stomach and endometrial cancers. We found that all three cancer types had the first two Gaussian clusters which consisted of low and high TMB samples, respectively. In colorectal and endometrial cancers, there was a third Gaussian cluster, in which samples possessed extremely high TMB. We named these three “hidden” subtypes as TMB-low, TMB-high and TMB-extreme. We further classified each sample to these three subtypes by Gaussian Mixture Model (GMM) for investigation of their significance.

The hypermutated phenotype can be caused by mutated *POLE* or MMR system defects. To gain insights on which mechanism may be responsible for the distinct TMB levels among three subtypes, we examined the non-synonymous mutations in *POLE* and seven MMR genes, and MSI status detected as described in earlier works^23–25^. We found that almost all of the TMB-high samples, 94%, 78% and 91% in colorectal, endometrial and stomach cancers, respectively, were MSI-High (MSI-H). A large fraction (92%) of TMB-extreme samples possessed at least one non-synonymous mutation in *POLE* in both colorectal and endometrial cancers. We observed relatively fewer MSI-H cases in the TMB-extreme subtype and fewer mutated *POLE* cases in the TMB-high subtype (Fig. 3). This can be possibly due to the mutually exclusive mechanisms for genomic instability. In previous studies^28^, MMR system defects have been linked to increased deletion/insertion (INDEL), which led us to explore INDEL rates among subtypes. We found that TMB-high samples generally had a significantly higher fraction (~17%) of INDEL mutations in contrast to what we observed in both TMB-low (~5%) and TMB-extreme (~1%) samples (Fig. 3). These distinct mutation profiles suggested that MMR system defects (MSI-H phenotype) were the likely cause for TMB-high and mutated *POLE* system for TMB-extreme.

**Fig. 3.**
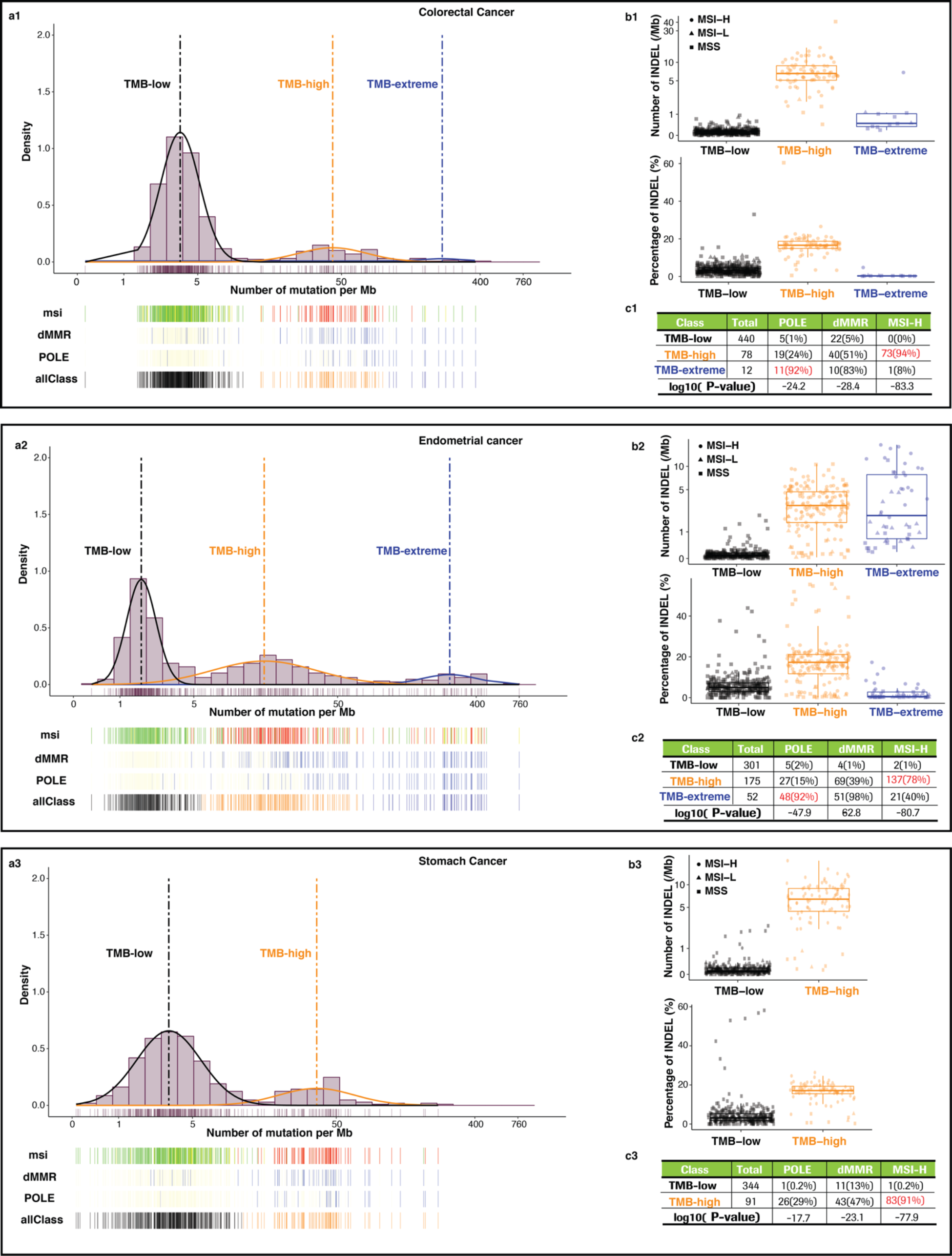
Three hidden subtypes revealed by log transformed TMB in colorectal, endometrial and stomach cancer. (a1-3) Distribution plots of log-transformed TMB for 1) colorectal, 2) endometrial, 3) stomach cancers. Three subtypes were determined by Gaussian Mixture Model classification and labeled with black (TMB-Low), orange (TMB-High) and blue (TMB-Extreme) in bar named allClass. MSI status for each subject is shown with green (MSS) and red (MSI-H) in msi bar. Non-synonymous mutation existence (occurrence > 1) in *POLE* or dMMR pathway genes, including *MLH1, MLH3, MSH2, MSH3, MSH6, PMS1, PMS2* are shown in blue and wild type is shown in yellow. (b1-3) Boxplots show INDEL mutation rates and percentages for three subtypes. (c1-3) Tables show total number of samples in each subtype, number of samples with at least one non-synonymous mutation in dMMR/*POLE* genes and number of samples with MSI-H status. The percentages of mutated *POLE*/MMR or MSI-H samples in each subtype are labeled in parentheses. Two-sided fisher exact tests were conducted to generate the p-value for each mutation profile among the subtypes.

Not all mutations have a deleterious impact on protein function. Therefore, we further investigated whether any driver mutations existed for TMB-high and TMB-extreme phenotypes. We compared non-synonymous mutations in POLE of TMB-extreme samples against the rest and compared non-synonymous mutations in seven MMR genes of TMB-high samples against the rest, using aggregated colorectal, stomach and endometrial cancer data (Fig. 4 and Supplementary Fig. 8). As expected, several driver mutations were discovered, including P286R and V411L in POLE, N674lfs*6 in MLH3 and K383Rfs*32 in MSH3 (Fig. 4). P286R and V411L in POLE, which were significantly enriched (binomial test p-values 1.38 * 10^−11^, and 5.88 * 10^−5^, respectively), were known driver mutations for the hypermutated phenotype^7^. N674lfs*6 in MLH3 and K383Rfs*32 in MSH3 have been detected in other studies^29,30^ but never reported as driver mutations for either MSI-H or hypermutation phenotypes. In this study, we found 10 out of 25 MSH3-mutated TMB-high samples had N674lfs*6 mutation in contrast to 0 out of 35 MSH3-mutated samples in TMB-low plus TMB-extreme subtype (p-value = 0). Additionally, 15 out of 36 TMB-high MSH3 mutated samples had K383Rfs*32 mutation in comparison with 1 out of 38 MSH3-mutated samples in TMB-low plus TMB-extreme subtype (p-value = 6.63 * 10^−15^). The high occurrence of these mutations in the TMB-high subtype suggested their potential driver effects in terms of resulting into MSI-H and relatively high TMB phenotypes.

**Fig. 4.**
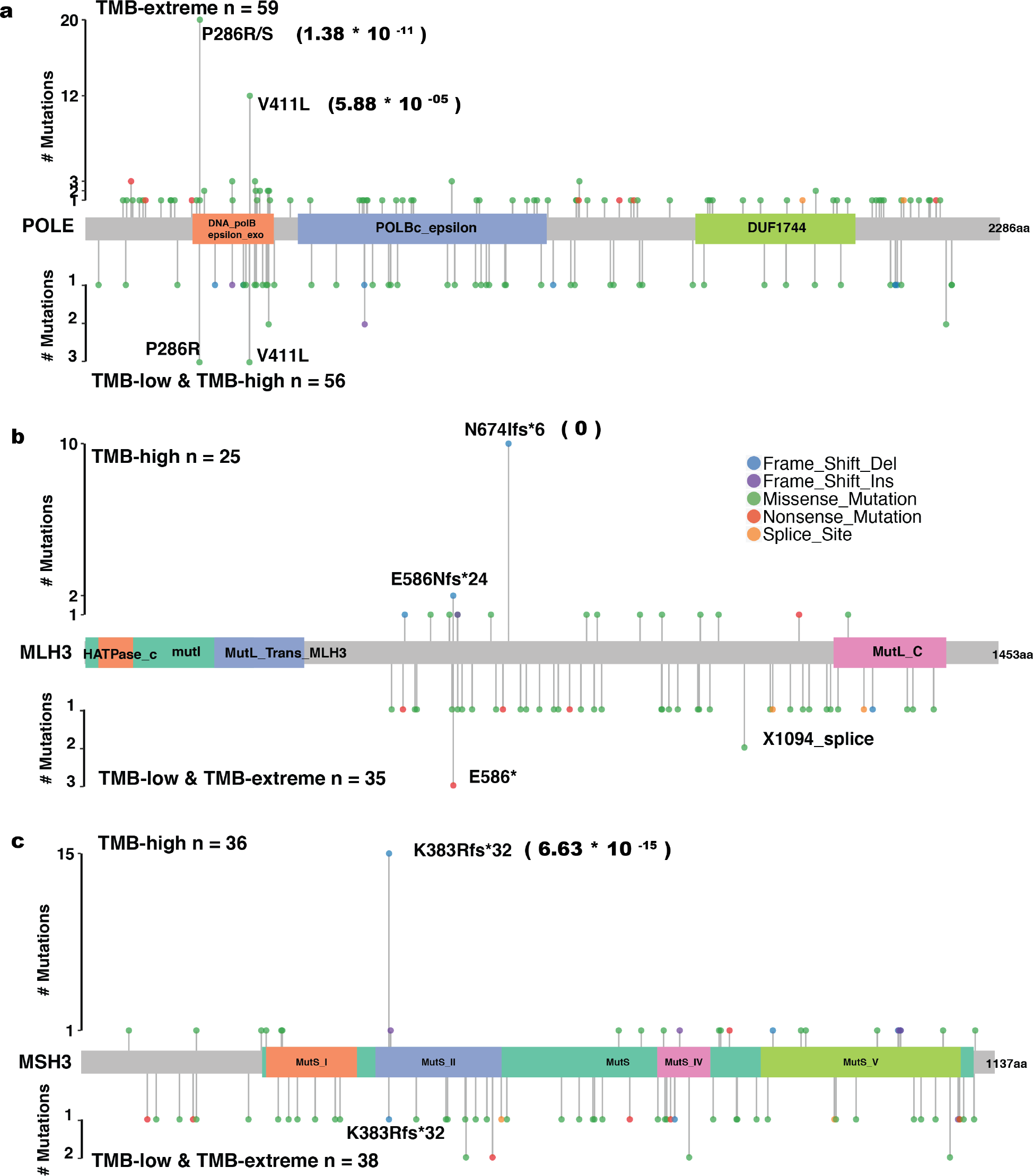
Known and novel *POLE*, *MLH3* and *MSH3* driver mutations. (a) Landscape of driver mutations in *POLE* detected in TMB-extreme group (top) compared with aggregated TMB-high and TMB-low group (bottom). Enrichment p-values using two-sided binomial test are shown in parentheses. (b-c) Landscape of driver mutations in *MLH3* and *MSH3* detected in TMB-high group (top) compared with aggregated TMB-extreme and TMB-low group (bottom). Enrichment p-values using binomial test is shown in parentheses.

To investigate the clinical relevance of the three subtypes derived by log-transformed TMB, we examined the associations of subtypes with tumor infiltrating immune cell abundance and overall patient survival. By using the immune infiltrates estimation for TCGA samples generated from Li T. et al study^31^, we found TMB-high and TMB-extreme samples had higher abundances of infiltrating CD8 T cell and Dendritic cell (DC) (Supplementary Fig. 9). The presence of cytotoxic CD8+ T cells, B cell and mature activated DCs in tumor microenvironment has been associated with a good clinical outcome in most cancer types^32^. We conducted survival analysis on aggregated colorectal, stomach and endometrial cancers because of the small size of the TMB-extreme group in colorectal cancer. We found that TMB-high and TMB-extreme were associated with improved patient survival at different levels after considering age and cancer stage (hazard ratio (HR) for TMB-high = 0.8 with p-value = 0.1; hazard ratio (HR) for TMB-extreme = 0.32 with p-value = 0.006) (Fig. 5), suggesting TMB subtypes are clinically relevant.

**Fig. 5.**
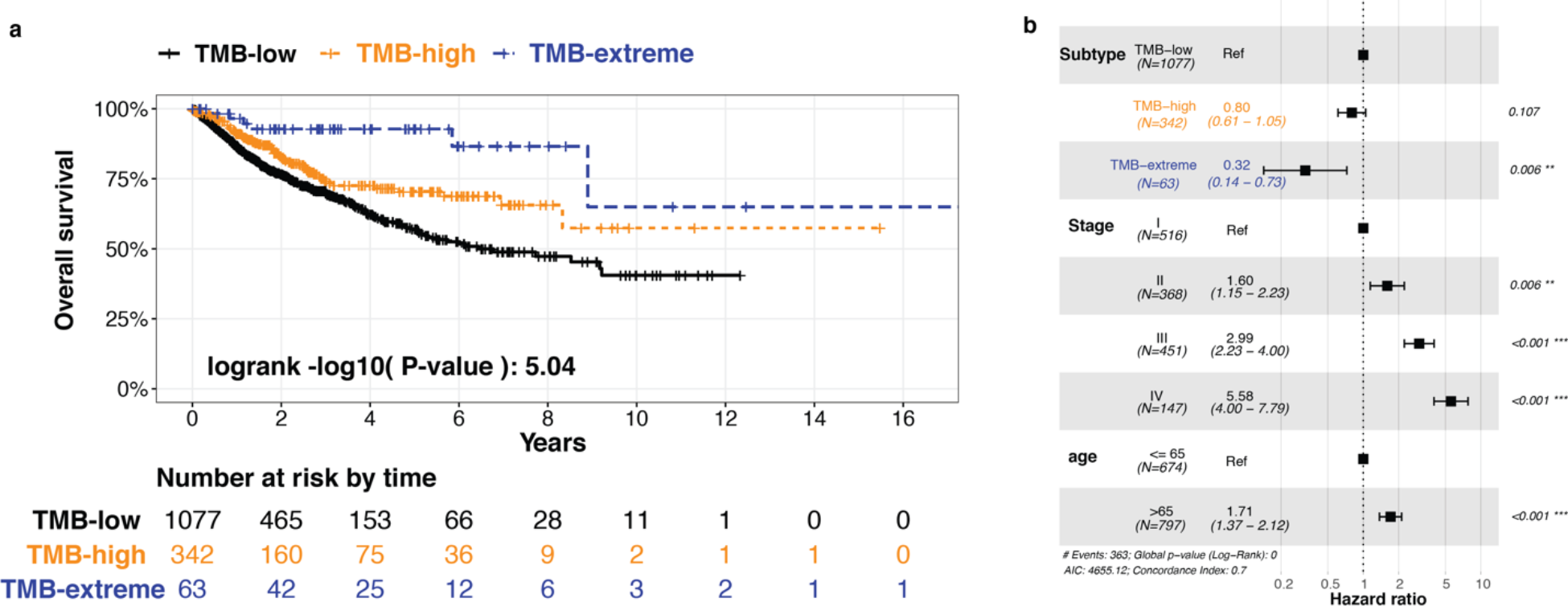
Survival outcome association with three subtypes. (a) Kaplan-Meier survival curves stratified by TMB subtypes using aggregated colorectal, endometrial and stomach patients. (b) Proportional hazard ratio analysis by Cox proportional-hazards model.

### Classification performance

With the discovery of biologically and clinically meaningful subtypes defined by log-transformed TMB, we extended the capability of our method to classify TMB subtypes using GMM (Fig. 1). Using the subtypes determined by WES-based TMB as truth, we evaluated the classification accuracy using panel-based TMB predicted by ecTMB and the counting method in the test sets. Comparing against the counting method, classification using ecTMB improved not only the overall accuracy and kappa concordance score, but also F1 score for each subtype classification (Fig. 6).

**Fig. 6.**
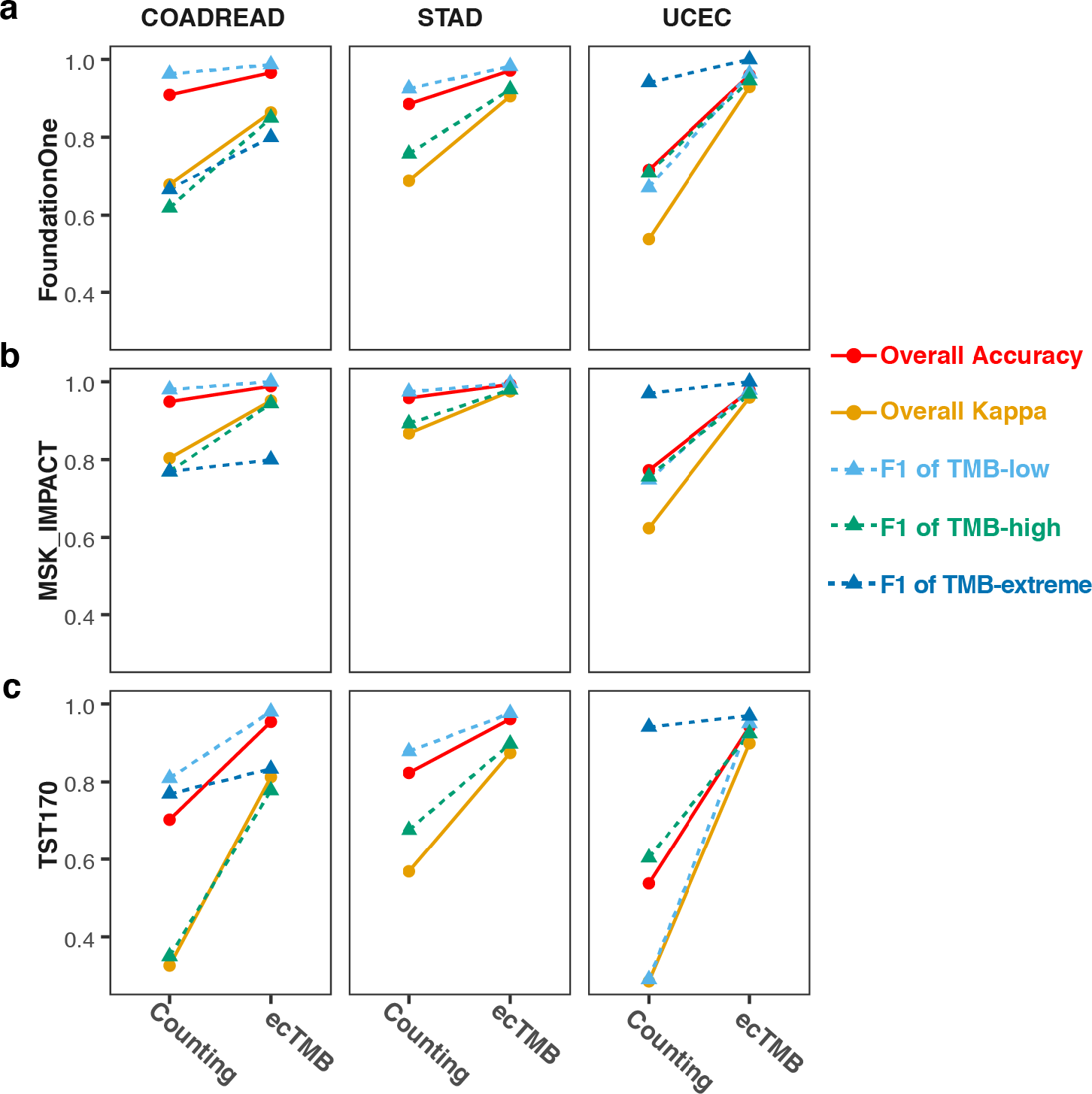
Comparison of TMB subtypes classification performance. (a-c) The plots show the comparison of overall accuracy (red), overall kappa score (orange) and F1 score for each subtype (TMB-low in cyan, TMB-high in green and TMB-extreme in blue) for TMB subtype classification using TMB predicted by ecTMB or counting method.

## Discussion

TMB is considered as the measure of neo-antigens in a tumor since it is historically calculated by number of non-synonymous mutations per Mb. However, another intuitive understanding of TMB is its equivalent to sample-specific BMR since the majority of mutations are passenger mutations in the whole exome. Thus, based on this second observation, we are the first to implement an explicit background mutation model for TMB prediction. Our background mutation model takes account of unknown as well as known mutational heterogeneous factors, including tri-nucleotide context, sample mutational burden, gene expression level, and replication timing by utilization of a Bayesian framework. We have shown that the three-step method successfully predicted background mutations, bringing several well-known cancer-specific driver genes to light. Compared to the counting method, ecTMB has several advantages. Firstly, ecTMB improves the consistency of TMB prediction among assays. Even though counting method predictions are highly correlated with WES-based TMB (Supplementary Fig. 11), the fixed or proportional biases cause the inconsistencies among assays. However, ecTMB is able to predict consistent TMB values in closer agreement with WES-based TMB in spite of the panel used, whether synonymous mutations are incorporated, or the proportion of non-synonymous mutations used as shown in this study. Secondly, ecTMB enables the integration of synonymous mutations for TMB prediction. Panel-targeted sequencing is desirable in clinical practice due to lower costs and fewer DNA input requirements but a reduced number of mutations per patient will be detected. The integration of synonymous mutations has the potential to improve the accuracy of panel-based TMB prediction. Lastly but not the least, ecTMB predicts TMB by considering each gene as an independent negative binomial process, which provides a more robust prediction compared to predicting TMB based on a single counting value. Although there are other factors influencing the consistency of TMB among assays, such as sequencing depth and choice of somatic mutation caller, we have demonstrated that ecTMB can help to improve the stability of TMB measurement when those factors are fixed. It is possible that more factors can be added to our statistical framework to further improve consistency of TMB measurements.

Different arbitrary cutoffs for TMB have been used to assess the biological and clinical interpretation of TMB subtypes. For example, some studies found an association between MSI-H and high TMB, wherein MSI-H is largely a subset^16^. However, there are no conclusive thresholds to define meaningful TMB subtypes. In our work, we discovered three cancer subtypes simply based on a log-transformed TMB, namely: TMB-low, TMB-high, and TMB-extreme. We have shown that these subtypes not only describe different levels of TMB but also are linked with various causes of hypermutation and overall patient survival. The first subtype is TMB-low, which has low mutation rate and very few mutations in *POLE* or MMR defects (MSI-H). The second subtype (TMB-high) is characterized with relatively high TMB, high INDEL mutation rate and high enrichment of MSI-H cases. This subtype is the subset that suffers from MMR system defects leading to MSI-H and relatively high TMB phenotype. Interestingly, we discovered two novel driver mutations for MMR defects, which need further research and clinical follow-up. The final subtype is TMB-extreme which is characterized by an extremely high SNV mutation rate but a low INDEL mutation rate, few MMR defects, and mutated *POLE*, where two known *POLE* driver mutations were discovered. This suggests that dysfunctional *POLE* might be the root cause of the TMB-extreme subtype. In all, our work is the first to clearly illustrate the association between MSI-H and high TMB, in which MSI-H is one subtype of hypermutated tumor. The novel TMB-extreme subtype shows even better overall survival outcomes compared to the TMB-high (MSI-H), suggesting that TMB-extreme might be another promising biomarker to predict patient prognosis or guide cancer treatment. The discovery of three TMB subtypes enabled us to extend ecTMB to classify samples based on predicted TMB values with a Gaussian Mixture Model.

However, we only detected these three distinct subtypes in colorectal, stomach and endometrial cancers, which are known to have a high percentage of MSI-H patients and other cancer types have very few MSI-H cases^33^. Therefore, these subtypes may be unique to cancers with a high percentage of MSI-H cases. Among other cancer types, we found that the majority of them have their own basic mutation rate represented by the first Gaussian (Supplementary Fig. 7), which might associate with their tissue type. For example, low grade glioma (LGG) has a lower basic mutation rate than esophageal carcinoma (ESCA) (Supplementary Fig. 7), which might be due to a lower rate of cell proliferation in brain than esophageal tissue. Cancers influenced with environmental factors have a continuous, broader spectrum of high TMB. Meanwhile, we detected hypermutated samples in these cancer types, which are also characterized by high mutation in *POLE* and MMR system, suggesting that a combination of other mutational biomarkers may help further classify these cancers.

In our studies, we described a powerful and flexible statistical framework for not only predicting accurate and consistent TMB measurements for various assays but also classifying samples to TMB subtypes which are biologically and clinically relevant. It presents another interpretation of TMB, which is sample-specific BMR, and sheds light on clinically relevant TMB subtypes. We believe that our method can help facilitate the adoption of TMB as biomarker in clinical diagnostics.

## Methods

### Datasets

Somatic mutations reported by MuTect2 (using the human genome reference build hg38) and clinical profiles of TCGA samples were downloaded from NCI Genomic Data Commons^34^ using TCGAbiolinks^35^. Formalin-fixed paraffin-embedded (FFPE) tissue samples were excluded from downstream analysis. The main cancer types included in our analyses were colon adenocarcinoma (COAD), rectal adenocarcinoma (READ), stomach adenocarcinoma (STAD), and uterine corpus endometrioid carcinoma (UCEC). Based on previous analysis, READ and COAD are often combined for analysis due to their similarity^23^. The availability of MSI status of these cancer types provided us an opportunity to investigate the association between TMB and MSI status. Sample IDs are listed in Supplementary Table 1. Totally, our study used 530 colorectal cancer samples (353 as training and 177 as test), 528 endometrial cancer samples (352 as training and 176 as test), and 436 stomach cancer samples (291 as training and 145 as test).

Tumor-infiltrating immune cell abundance data was downloaded from TIMER^31^. Replication timing, expression level, and open-chromatin state of all protein-coding genes were extracted from Lawrence et al.^3^.

### Whole exome annotation

Ensembl build 81 (for the GRC38 human reference) was downloaded and processed to generate all possible mutations and their functional impacts for the genome. First, we changed every genomic base in coding regions to the other three possible nucleotides and used Variant Effect Predictor (VEP) to annotate their functional impacts. Each variant’s functional impact was chosen based on the following criteria: biotype > consequence > transcript length. Each variant’s tri-nucleotide contexts, including before and after the mutated base, and corresponding amino acid positions relative to protein length were reported.

### TMB prediction method

Our proposed novel TMB prediction method adapted background mutation modeling and added a final step to predict TMB using maximum likelihood for a given new sample. A training cohort with whole-exome sequencing can be used to estimate parameters of BMR model. The modeling steps of TMB prediction are described below.

#### 1. Mutation rate for each sample (b_s_)

The mutation rate for each sample (b_s_) can be determined by the total number of mutations of the sample divided by size of evaluated genome in Mb (Megabase) unit. If only non-synonymous mutations were used, b_s_ is equivalent to current standard TMB calculation. Here, we used the number of non-synonymous mutation for the following TMB prediction and classification.

#### 2. tri-nucleotide context-specific mutation rates

Tri-nucleotide context-specific mutation rates were estimated for the training cohort. We considered the 96 possible tri-nucleotide contexts (from the 6 possible types of single base substitutions - A/T->G/C, T/A->G/C, A/T->C/G, T/A->C/G, A/T->T/A, G/C->C/G - and possible nucleotides around it, plus indels. Mutations are classified as synonymous or non-synonymous based on whether they cause a change to the amino acid sequence of the translated protein. We assume that synonymous mutations occur according to the BMR.

For each tri-nucleotide mutation context *i*, we calculate the number of synonymous *n*_*i*(*synonymous*)_ and non-synonymous *n*_*i*(*nonsynonymous*)_ mutations observed across all tumor samples and the number of possible synonymous *n*_*i*(*synonymous*)_ and non-synonymous *n*_*i*(*nonsynonymous*)_ variants in the exome. For non-synonymous mutations, only genes that are less likely to be drivers, are taken into consideration to avoid skewing the background non-synonymous mutation rate; i.e., the bottom 60% of genes ranked by the number of mutated samples in decreasing order. The potential bias introduced by using a subset of genes for non-synonymous mutations is corrected by the factor *r*, which is estimated using the method of moment^36^, calculated as the mean of 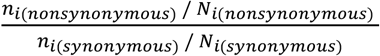 across all mutation contexts. For mutation context *i*, the mutation rate *m*_*i*_ is calculated as shown in Equation [1]. When calculating indel mutation rate *m*_*indel*_, we assume that all protein-coding positions can have indels, and that all indels are considered as non-synonymous.

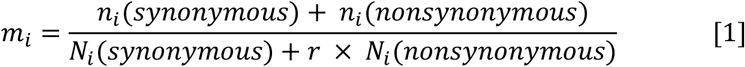

#### 3. Gene-specific mutation rate *α*_*g*_

##### 3.1 Regression model across genes

We assume that the occurrence of synonymous mutations represents the background mutation rate, and we can model the number of synonymous mutations per gene using the negative binomial.

We consider multiple factors that could influence the underlying mutation rate to model the synonymous mutation counts. Firstly, the number of possible synonymous mutations is controlled by the gene’s coding sequence (e.g. codons and length). Specifically, for gene *g*, we get all possible bases that could mutate to synonymous mutations and sum up their context-specific mutation rate as *E*_*g*(*synonymous*)_ = *∑*_*synonymous*_ *m*_*i*_. Secondly, because different individuals are expected to have different background mutation rates, we use the aforementioned sample specific factor *b*_*s*_ to represent the total mutation burden of sample *s*. Thirdly, *α*_*g*_ is gene-specific mutation rate, influenced by several additional known factors that can influence the underlying mutation rate for a given gene, including replication timing (R), expression level (X), open-chromatin state (C), and whether gene is an olfactory receptor (O). The effect of these factors is estimated using negative binomial regressions as described below.

We model the synonymous mutation count *y*_*gs*_ of gene *g* and sample *s* with negative binomial regression [2], assuming common dispersion *p* across genes.

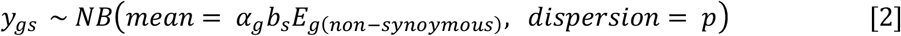

Where

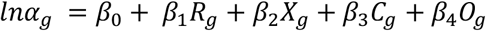

*β* = {*β*_0_, *β*_1_, *β*_2_, *β*_3_, *β*_4_} is estimated by performing regression with all genes.

##### 3.3 Capture unknown factor impact through Maximum Likelihood Method

In [2] we assume mutation rate factors only dependent on proposed regressors, but unknown mechanisms or biological factors can also impact mutation rate. Therefore, we model each gene as an independent zero-inflated Poisson process and use the Maximum Likelihood estimation (MLE) to estimate gene-specific dispersion parameter 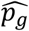 and gene-specific mutation rate 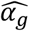 by maximizing [3].

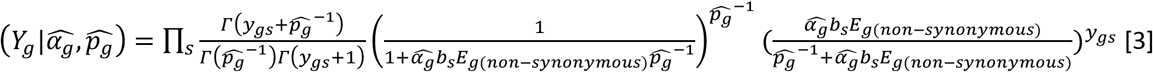

For each gene, *Y*_*g*_ = {*y*_*g*1_, *y*_*g*2_, …, *y*_*gs*_} are the silent mutation counts in different samples, the initial value of 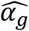 is given by 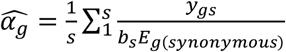, and the influencing factors (*R*, *X*, *C*, *O*) are not applicable.

##### 3.4 Optimization of gene-specific mutation rate factors

Because we obtain *α*_*g*_ by pooling all genes together, it captures the common trend of the influencing factors (*R*, *X*, *C*, *O*) on background mutation rate. On the contrary, 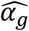 is a gene-specific parameter from the observed data independent of the influencing factors. 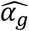 and *α*_*g*_ are not always the same, which could be caused by technical noise (e.g. errors in mutation calling algorithms) or reflect real biological mechanisms (e.g. factors influencing the background mutation rate that are not included in our regression model). Due to the low number of somatic mutations in each gene, 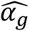 is very vulnerable to statistical noise. Thus, we try to find the optimized 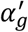 by incorporating both parameters from negative binomial regression and those directly from gene-specific estimation. The posterior probability of 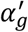 is proportional to the likelihood times prior with *σ* estimated as [7]. The prior probability is chosen to constrain 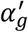 to be centered at *α*_*g*_. We maximize [4] to obtain the proper 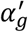 for each gene.

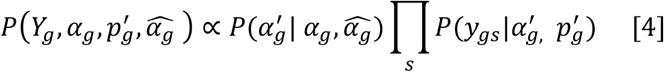

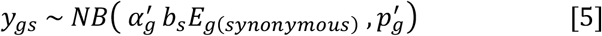

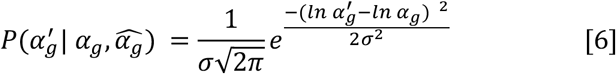

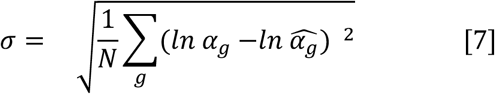

Then the ‘gene-specific estimation’ and ‘optimization of gene mean’ steps are repeated by replacing 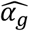 with 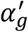 to re-estimate dispersions until convergence is achieved. The estimated 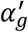 and 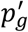 are used in the following steps.

#### 4. TMB prediction

Using pre-defined 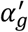 and *p*_*g*_, we modeled each gene as an independent negative binomial process for a given new sample s’. Then, we use Maximum Likelihood estimation (MLE) to estimate *b*_*s*_ by maximizing [8]. In this step, *Y*_*g*_ = {*y*_1_, *y*_2_, …, *y*_*g*_} captures the synonymous mutation counts, and additionally, non-synonymous mutation counts in sample s’. When WES is used, *Y*_*g*_ includes all gene in the genome. When particular panel is used, *Y*_*g*_includes the genes covered by the panel.

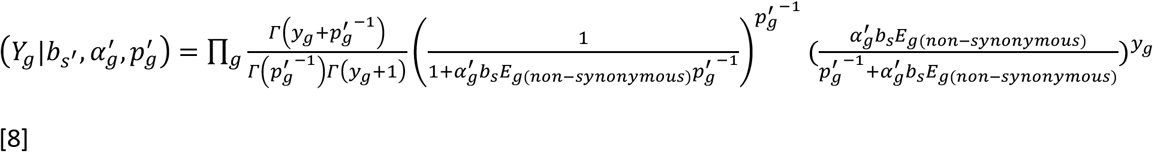

Poisson and zero-inflated Poisson regressions can also be used for background mutation modeling which is also implemented in the package. We used negative binomial for all analyses presented in this study.

### TMB prediction evaluation

The TMB varied in a large range from 0.01 per Mb to 760 per Mb and the majority of samples (76%) had TMB less than 10 per Mb. Therefore, we presented performance measures with log-transformed values along with non-log-transformed ones to deal with the large dynamic range of data and to avoid the mean absolute error being determined only by large numbers. In order to avoid dealing with zeros, TMB values were incremented by one before log-transformation. MAE, RMSE and Bland-Altman analyses were conducted on log-transformed values and non-log-transformed values. Bland-Altman analysis is a widely used method to assess the agreement between two different assays, providing a bias measurement (mean difference), the limits of agreement and 95% confidence intervals for these measurements. Bland-Altman analysis was done using the R package blandr.

### TMB subtype classification

Log-transformed TMBs were modeled using Gaussian Mixture Model [9], in which components represent cancer subtypes. The Expectation-Maximization algorithm can be used to estimate each component’s parameters in the Gaussian mixture model with training data. The parameters for K^th^ component include weight (π^*k*^), mean (*μ*^*k*^), and variance (*∑*^*k*^). These parameters are used in assignment score calculation.

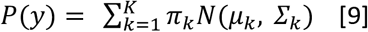

For a given new sample’s log-transformed TMB (*y*_i_), assignment score for each component (*γ*(*b*|*C*_*k*_)) will be calculated as [10] using pre-defined parameters.

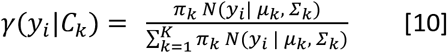

The new sample will be classified to the component which has the highest assignment score. Currently, weight = 1 was assigned to each component due to under-representation of TMB high samples.

### Cluster cancer types based on TMB distribution

WES mutation data for 29 cancer types other than colorectal, endometrial and stomach cancers were downloaded from GDC. However, many cancer types, such as adrenocortical carcinoma (ACC), do not have a significant number of hypermutated samples. In order to have a large population of hypermutated samples, we considered aggregating cancer types together. However, we discovered that mutation spectra among cancer types were different as well, indicating a different threshold for hypermutated population for each cancer. For example, the median mutation rate of skin cutaneous melanoma (SKCM) is ~ 10 mutation per Mb and median of acute myeloid leukemia (LAML) is less than 1 mutation per Mb. Therefore, we decided to cluster cancer types based on the similarity of log-transformed TMB distribution (Supplementary Fig. 6) so that we check the distribution of log-transformed TMB within each group.

For each cancer type, the density of log-transformed TMB was generated by bin = 1. Then we used K-means clustering method to group cancer types to 5 clusters based on the similarity of the log-transformed TMB density. In each cluster, the mutation data is aggregated for further analysis.

### Cancer subtype classification and characterization

Within each cancer type (colorectal, endometrial and stomach cancer), log-transformed TMBs, either defined by total number of mutations per Mb or number of non-synonymous mutation per Mb, were modeled using Gaussian Mixture Model as described above. Each sample was assigned to one of TMB-low, TMB-high and TMB-Extreme classes based on its assignment score. For each sample, INDEL incidence, estimated immune cell abundance and non-synonymous mutation existence (occurrence > 1) in *POLE* and dMMR pathway genes including *MLH1, MLH3, MSH2, MSH3, MSH6, PMS1*, and *PMS2* were summarized. Mutations of *POLE* and MMR system genes were plotted using maftools^37^.

### Background mutation prediction by BMR model

Within each cancer type, WES data from training set was used to determine parameters for background mutation model either by simply GLM or three-steps approach. Background mutations were predicted based on [11] for non-synonymous mutation and [12] for synonymous mutation in both training set and rest of test set.

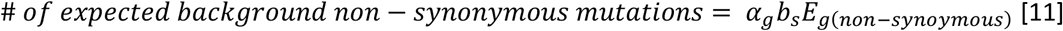

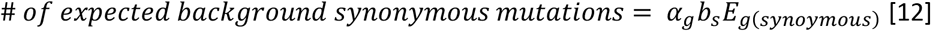

### Cancer survival analysis

Kaplan-Meier survival analysis was used to estimate the association of cancer subtype with the overall survival of patients with colorectal, endometrial and stomach cancers aggregated data. Furthermore, we performed proportional hazard ratio analysis using the coxph function in R, including age, stage and subtypes as covariates. The significances of the covariates were assessed by Wald tests. Overall survival was calculated from the date of initial diagnosis of cancer to disease-specific death (patients whose vital status is termed dead) and months to last follow-up (for patients who are alive).

### TMB prediction for panels

In order to evaluate ecTMB prediction for panels, we performed *in silico* analyses. We downloaded panel coordinates bed file of Illumina TruSight Tumor 170 (panel size 524kb) from Illumina website (https://support.illumina.com/content/dam/illumina-support/documents/downloads/productfiles/trusight/trusight-tumor-170/tst170-dna-targets.zip). The gene lists of FoundationOne CDx and Integrated Mutation Profiling of Actionable Cancer Targets (MSK-IMPACT) were download from the Foundation Medicine website (https://www.foundationmedicine.com/genomic-testing/foundation-one-cdx) and FDA (https://www.accessdata.fda.gov/cdrh_docs/reviews/den170058.pdf), respectively. Corresponding panel coordinates bed files were generated based on gene lists for FoundationOne CDx and MSK-IMPACT. Due to the lack of exact panel coordinates of FoundationOne CDx and MSK-IMPACT, the sizes of the panels converted from gene lists were larger than the real commercial panels. The final sizes of FoundationOne CDx and MSK-IMPACT panels were 5.4Mb and 10Mb, respectively. These bed files were listed in Supplementary Table 2.

Mutations located in a given panel were selected to represent the mutations which can be detected by targeted sequencing for that panel. Within each cancer type, WES data of training set were used to determine background mutation model parameters. *In silico* performance evaluations were done using the test data. Both ecTMB and counting methods were applied to test data.

## List of abbreviations

Mb: megabase
NGS: Next-Generation Sequencing
WES: whole-exome sequencing
TMB: tumor mutational burden
GLM: Generalized Linear Model
BMR: Background mutation rate
MUC16: membrane-associated mucin
TTN: titin
MLE: Maximum Likelihood Estimation
ACC: adrenocortical carcinoma
SKCM: skin cutaneous melanoma
LAML: acute myeloid leukemia
LUSC: lung squamous cell carcinoma
LUAD: lung adenocarcinoma
BLCA: bladder urothelial carcinoma
TILs: tumor infiltrating lymphocytes

## Declarations

### Acknowledgements

The results published here are largely based upon data generated by the TCGA Research Network (http://cancergenome.nih.gov/), and our use of these data is in accordance with the guidelines at: http://cancergenome.nih.gov/publications/publicationguidelines. We thank the TCGA Consortium for use of these datasets.

### Author contributions

L.Y., M.M., H.Y.K.L conceived the project. L.Y. and Y.F. designed the algorithm. L.Y. conducted the analyses and wrote the manuscript. M.M. and H.Y.K.L edited the manuscript. M.M. and H.Y.K.L supervised the work.

### Data availability

Data for this study was generated by TCGA, downloaded from NCI Genomic Data Commons (https://gdc.cancer.gov/). Tumor-infiltrating immune cell abundance data was downloaded from TIMER website (https://cistrome.shinyapps.io/timer/). Replication timing, expression level, and open-chromatin state of all protein-coding genes were extracted from Lawrence et al.^3^ (https://media.nature.com/original/nature-assets/nature/journal/v499/n7457/extref/nature12213-s2.xls). Illumina TruSight Tumor 170 panel coordinates bed file was downloaded from Illumina’s website (https://support.illumina.com/content/dam/illumina-support/documents/downloads/productfiles/trusight/trusight-tumor-170/tst170-dna-targets.zip) and the gene lists for FoundationOne CDx and MSK-IMPACT were downloaded from Foundation Medicine’s website (https://www.foundationmedicine.com/genomic-testing/foundation-one-cdx) and the FDA’s website (https://www.accessdata.fda.gov/cdrh_docs/reviews/den170058.pdf), respectively.

