## Supplemental figures for "ecTMB: a robust method to estimate and classify tumor mutational burden"

#### Supplementary Figures

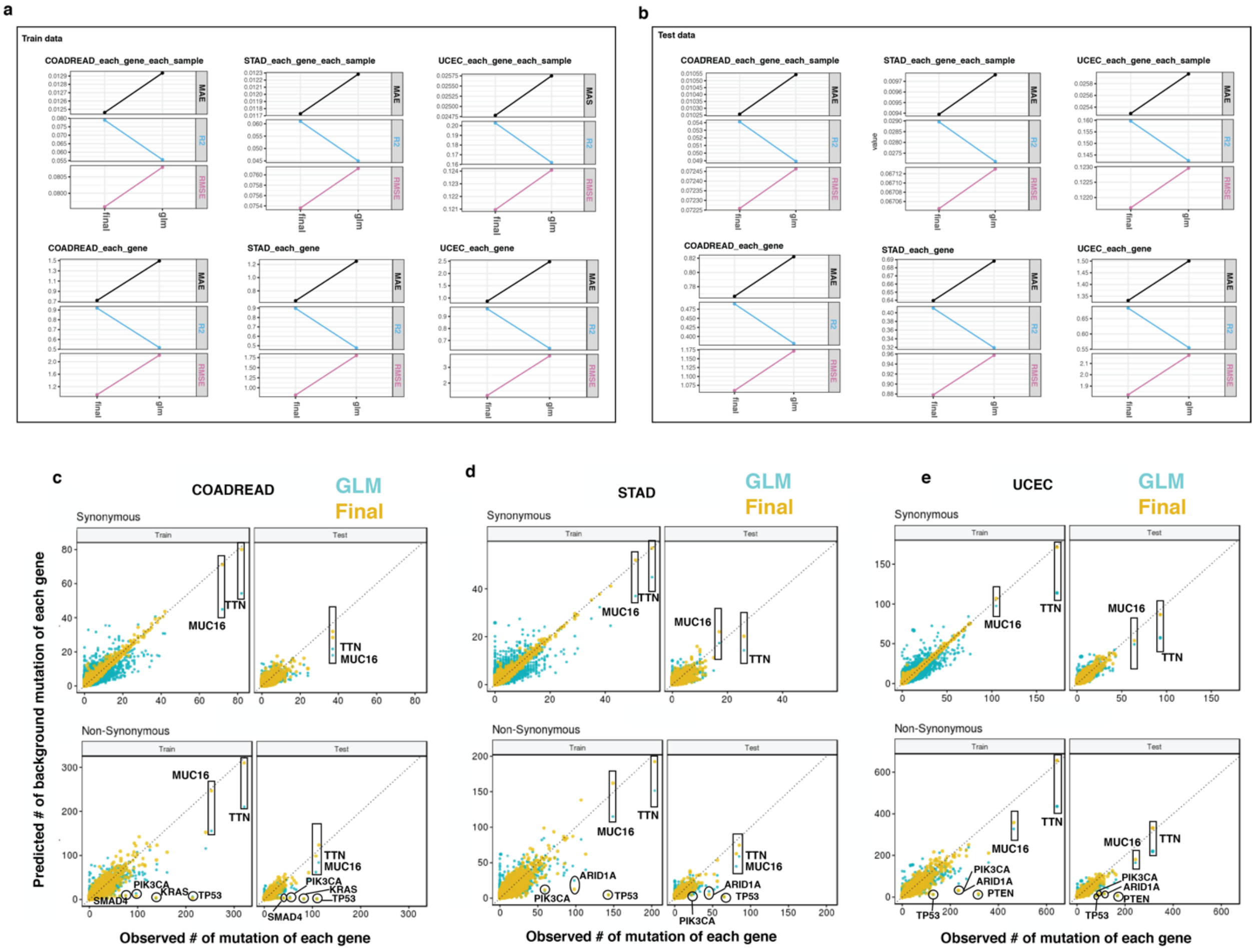


**Supplementary Fig. 1- Background mutation modeling comparison.** (a-b) Plot shows the comparisons of model accuracy between GLM model and final (3-steps) approach in training sets (a) and in test sets (b). RMSE, MAE and R-squared were calculated between predicted number of synonymous mutations and observed value for each gene in each sample (top) and each gene in aggregated samples (bottom). (c-e) Predicted number of background synonymous (top)/non-synonymous (bottom) mutation of each gene are plotted against observed ones in colorectal (c), stomach (d) and endometrial (e) cancers. The prediction made by GLM model is labeled in cyan and final (3-steps) approach in yellow. Several well-known driver genes are circled and labeled in the non-synonymous figures.

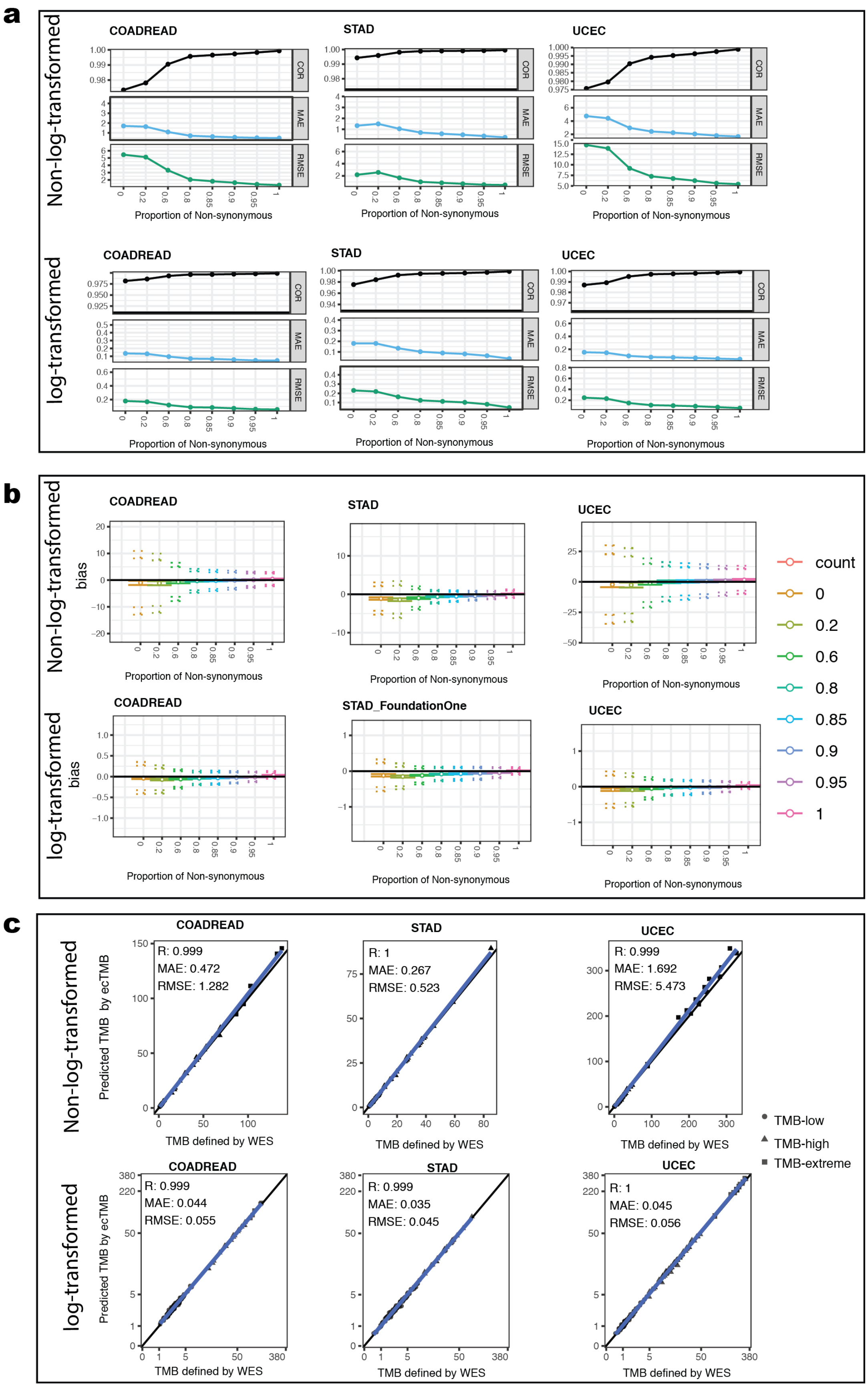
a
Non-log-transformed
log-transformed
COADREAD
STAD
UCEC
COR
MAE
RMSE
Proportion of Non-synonymous
b
bias
STAD_FoundationOne
count
0
0.2
0.6
0.8
0.85
0.9
0.95
1
c
Predicted TMB by ecTMB
TMB defined by WES
R: 0.999
MAE: 0.472
RMSE: 1.282
R: 1
MAE: 0.267
RMSE: 0.523
R: 0.999
MAE: 1.692
RMSE: 5.473
R: 0.999
MAE: 0.044
RMSE: 0.055
R: 0.999
MAE: 0.035
RMSE: 0.045
R: 1
MAE: 0.045
RMSE: 0.056
TMB-low
TMB-high
TMB-extreme

**Supplementary Fig. 2- ecTMB performance using WES data.** (a) Plot shows the comparisons of prediction accuracy when different proportions of non-synonymous mutation were used. RMSE, MAE and correlation coefficient were calculated between predicted TMB and standard WES-based TMB before log-transformation (top) and after log-transformation (bottom). (b) Biases, upper and lower limits are shown when various proportions of non-synonymous mutation were used. The results using non-log-transformation value are shown in top and log-transformation in bottom. The middle circle indicates the bias (mean difference) and two solid lines around it are 95% confidence interval for the bias. The two dotted line on the top are 95% confidence intervals for the upper limit of 95% agreement and ones on the bottom are 95% confidence intervals for the lower limit of 95% agreement. (c) Predicted TMB values are plotted against standard WES-based TMB before log-transformation (top) and after log-transformation (bottom). The linear regression lines were added. Standard WES-based TMB was calculated by counting number of non-synonymous mutation and then divided by size of the exome.

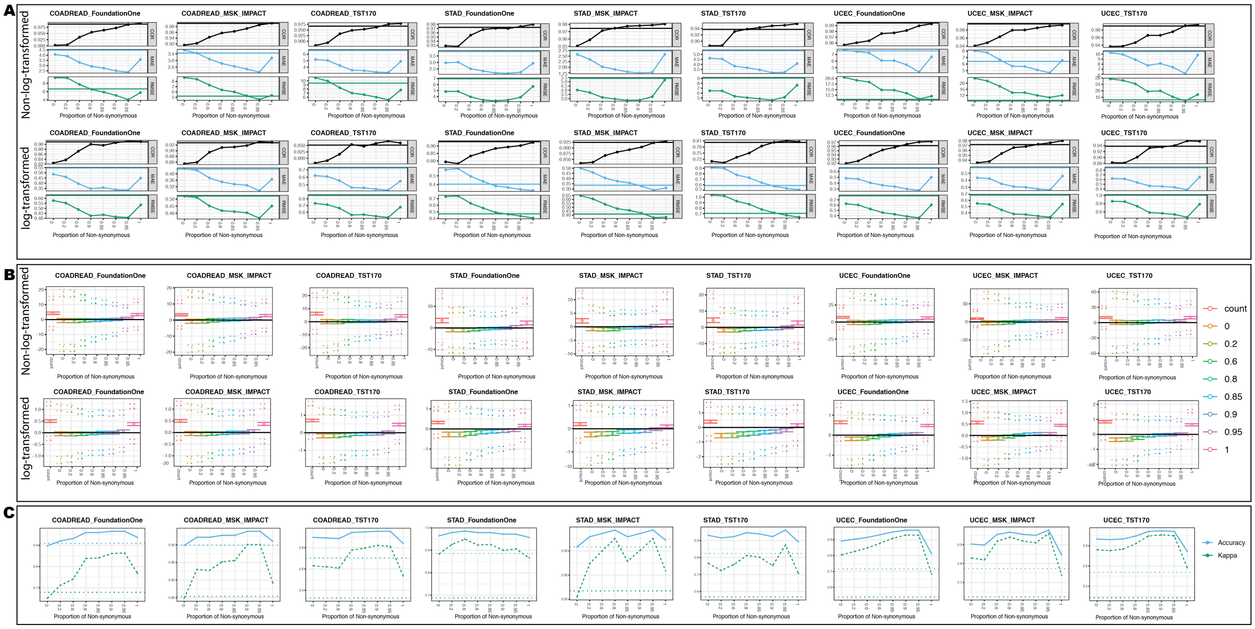


**Supplementary Fig. 3- Panel TMB prediction by ecTMB.** (a) Plot shows the comparisons of prediction accuracy when different proportions of non-synonymous mutation were used for each cancer and each panel. RMSE, MAE and correlation coefficient were calculated between predicted panel-based TMB and standard WES-based TMB before log-transformation (top) and after log-transformation (bottom). The horizontal line in each plot indicates the measurement when counting method was used, which simply counts the number of non-synonymous mutation per Mb. (b) Biases, upper and lower limits were presented when various proportions of non-synonymous mutation were used. The first column of each figure shows the Bland Altman analysis for TMB prediction by counting method. The results using non-log-transformed values are shown in top and log-transformation in bottom. The middle circle indicates the bias (mean difference) and two solid lines around it are 95% confidence interval for the bias. The two dotted lines on the top are 95% confidence intervals for the upper limit of 95% agreement and ones on the bottom are 95% confidence intervals for the lower limit of 95% agreement. The individual Bland Altman plot is shown in Supplementary Fig. 10. (c) Plots show the overall accuracy and kappa score for classifications of 3 TMB subtypes by ecTMB when different proportions of non-synonymous mutation were used. The horizontal dashed lines in each plot indicates the measurements when counting method was used.

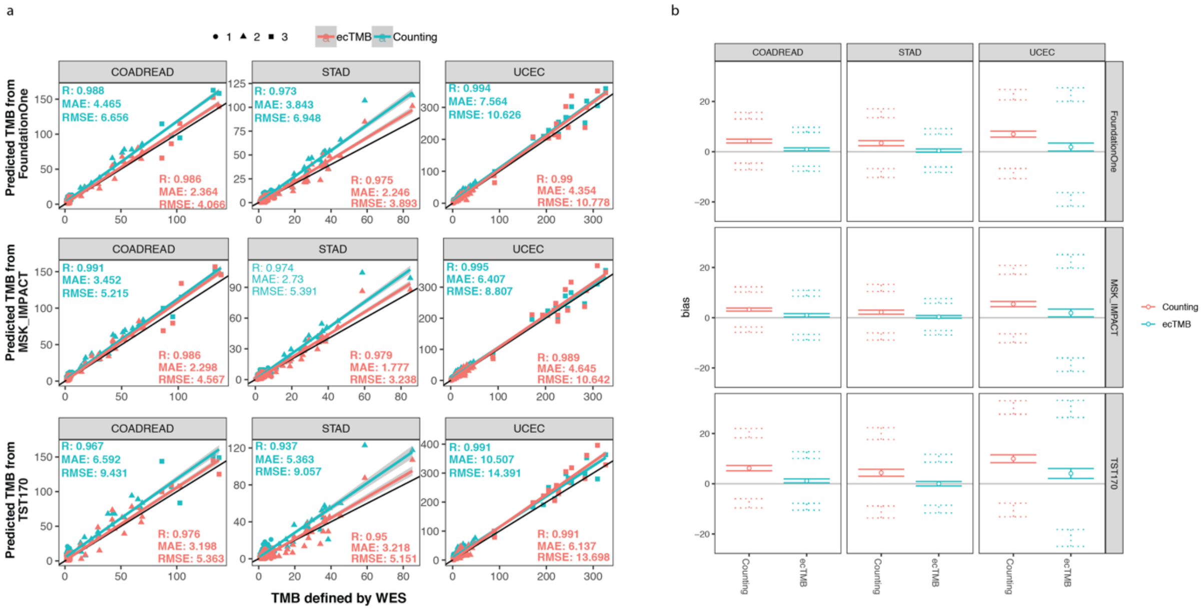


**Supplementary Fig. 4 – Evaluation of the panel-based TMB prediction performance before log-transformation.** (a) Scatter plots show WES-based standard TMB plotted against predicted panel-based TMBs for each cancer type and each panel. Two methods were used for panel-based TMB predictions, including counting method (in cyan) and ecTMB (in red). Their linear regression lines against WES-based TMB and performance measurements (correlation coefficient, MAE and RMSE) are plotted for each method in each scatter plot. (b) Bland Altman analysis results for counting method (cyan) and ecTMB (red) against WES-based TMB are shown here. The middle circle indicates the bias (mean difference) and two solid lines around it indicate the 95% confidence interval for the bias. The two dotted lines on the top are 95% confidence intervals for the upper limit of 95% agreement and ones on the bottom are 95% confidence intervals for the lower limit of 95% agreement.

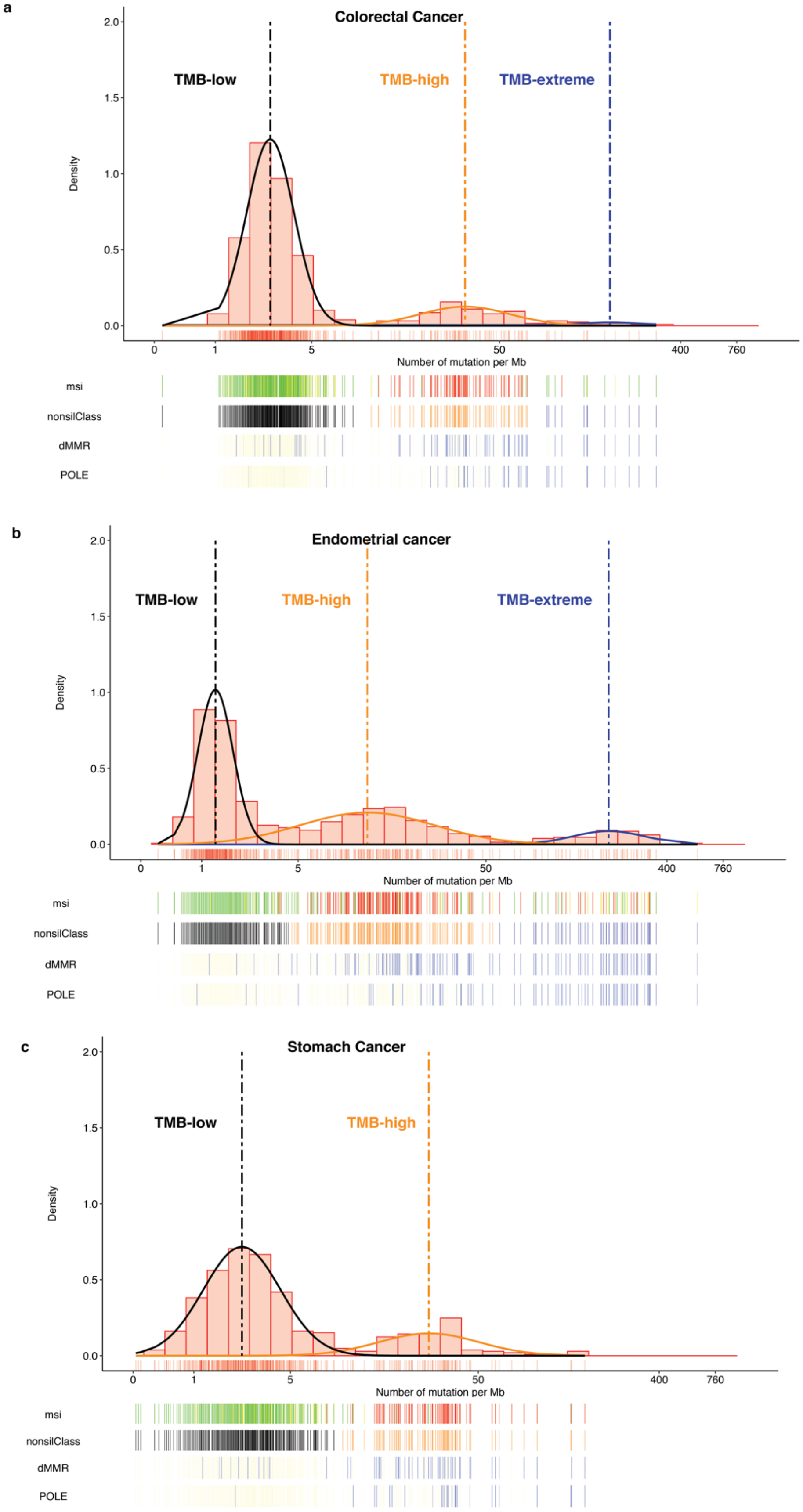
a
Colorectal Cancer
TMB-low
TMB-high
TMB-extreme
Density
2.0
1.5
1.0
0.5
0.0
0
1
5
50
400
760
Number of mutation per Mb
msi
nonsilClass
dMMR
POLE
b
Endometrial cancer
TMB-low
TMB-high
TMB-extreme
Density
Number of mutation per Mb
msi
nonsilClass
dMMR
POLE
c
Stomach Cancer
TMB-low
TMB-high
Density
Number of mutation per Mb
msi
nonsilClass
dMMR
POLE

**Supplementary Fig. 5 – Three hidden subtypes revealed by log transformed TMB calculated by number of non-synonymous mutation per Mb.** (a-c) Distribution plots of log transformed TMB for a) colorectal, b) endometrial, c) stomach cancers. Three subtypes were determined by Gaussian Mixture Model classification and labeled with black (TMB-Low), orange (TMB-High) and blue (TMB-Extreme) in allClass bar. MSI status for each subject is shown with green (MSS) and red (MSI-H) in msi bar. Non-synonymous mutation existence (occurrence > 1) in *POLE* or dMMR pathway genes, including *MLH1, MLH3, MSH2, MSH3, MSH6, PMS1, PMS2* are shown in blue and wild type is shown in yellow.

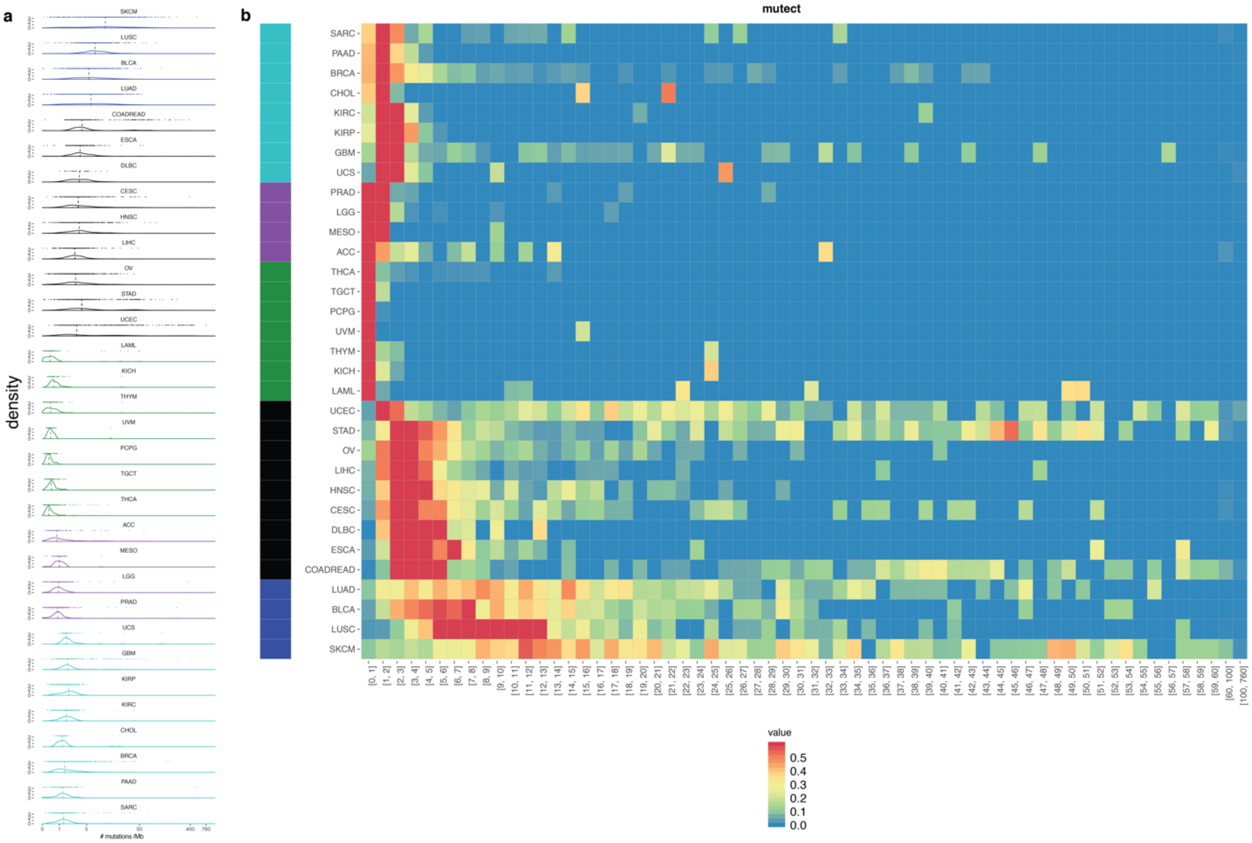


**Supplementary Fig. 6 – Cancer groups based on distribution of log-transformed TMB.** (a) Distribution plots of TMB for each cancer type in log scale. (b) Heatmap of distribution of log-transformed TMB. K-means clustering method was used to generate 5 clusters which is shown on the left side.

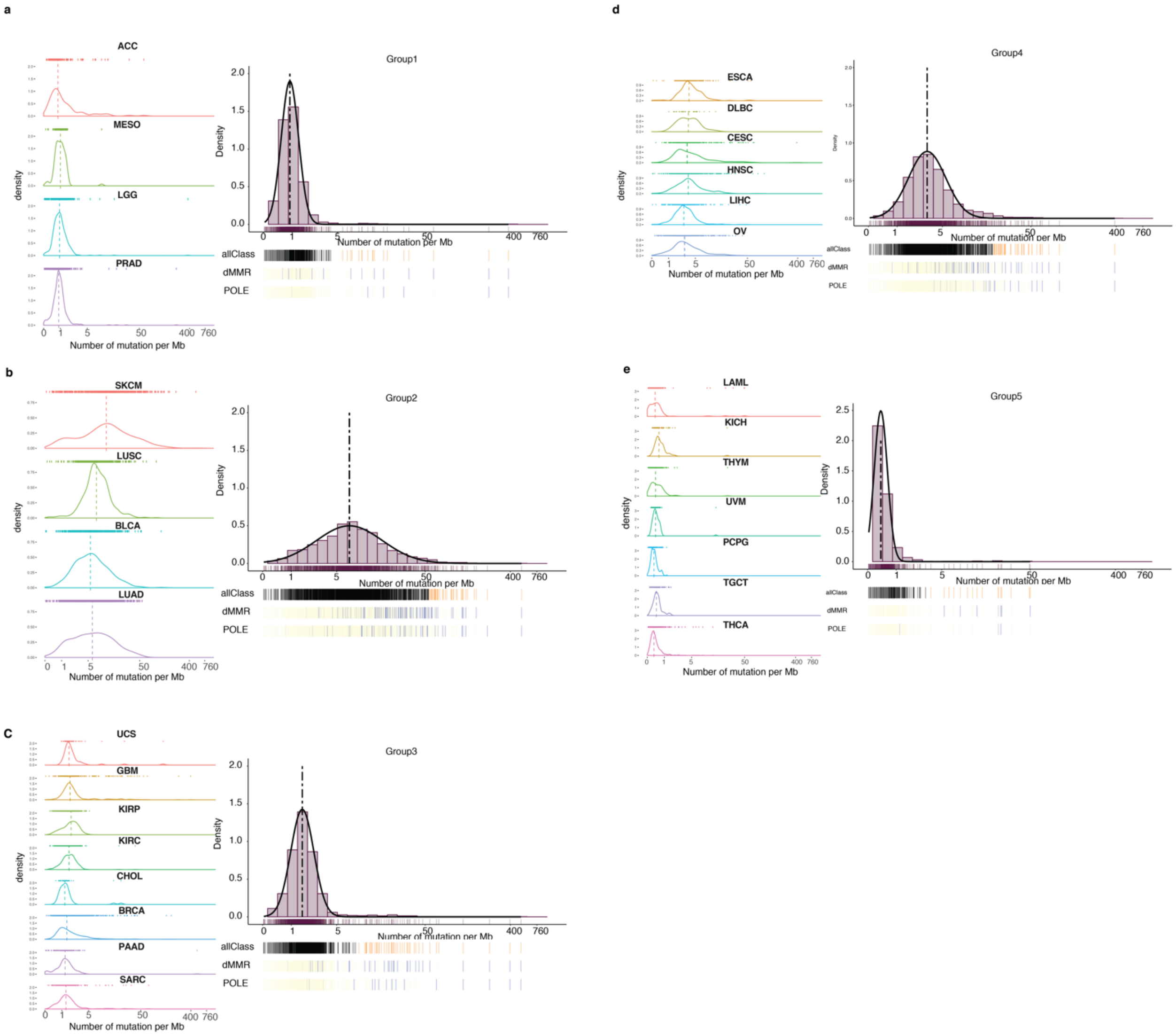


**Supplementary Fig. 7 –Distribution for each cancer group.** (a-e) Distribution of log-transformed TMB for each cancer: group 1 (a), group 2 (b), group 3 (c), group 4 (d) and group 5 (e). The distribution of log-transformed TMB for each individual cancer in each group is shown on the left.

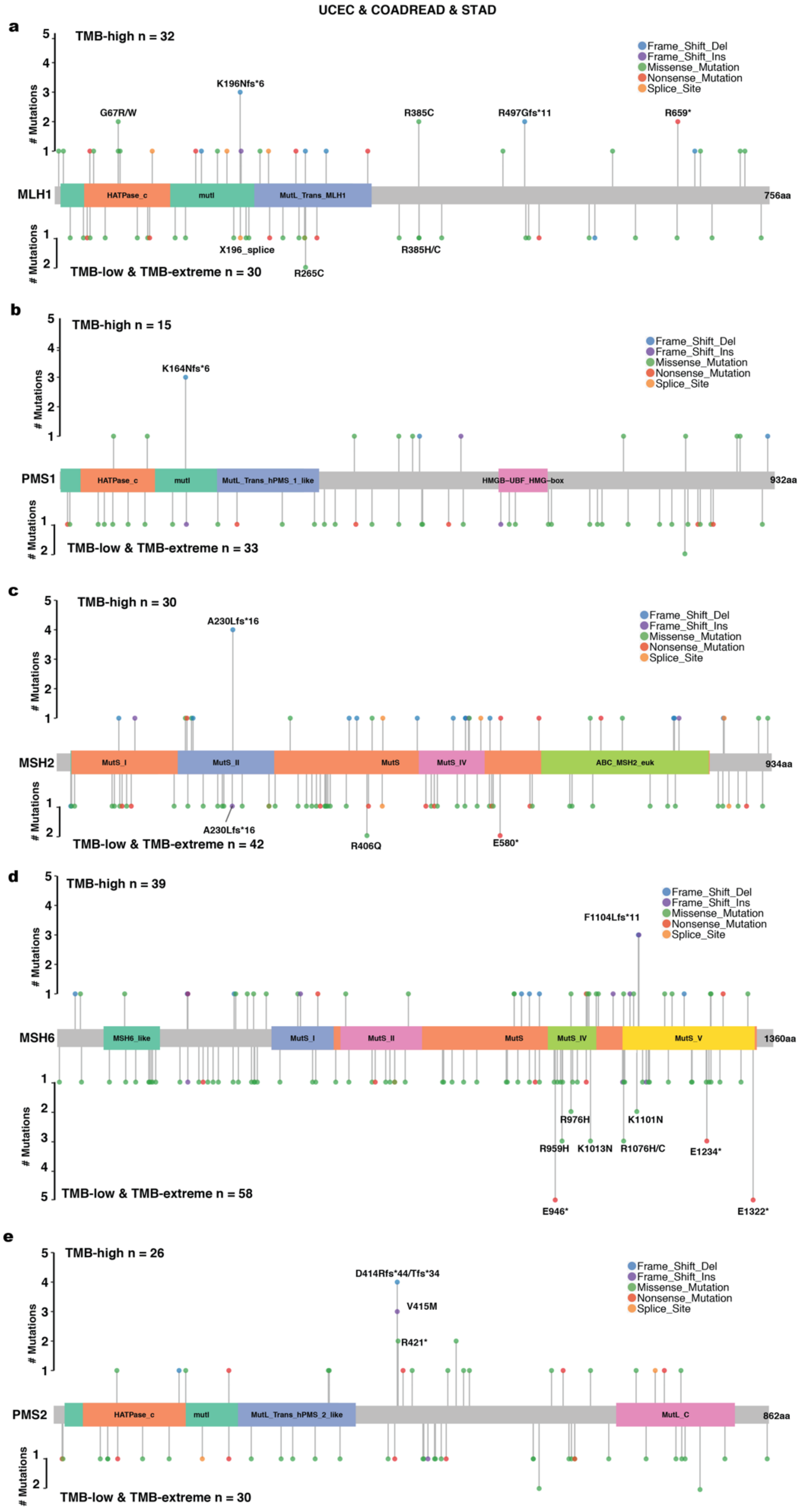
UCEC & COADREAD & STAD
a
TMB-high n = 32
### Mutations
Frame_Shift_Del
Frame_Shift_Ins
Missense_Mutation
Nonsense_Mutation
Splice_Site
K196Nfs*6
G67R/W
R385C
R497Gfs*11
R659*
MLH1
HATPase_c
mutl
MutL_Trans_MLH1
756aa
X196_splice
R385H/C
R265C
TMB-low & TMB-extreme n = 30
b
TMB-high n = 15
K164Nfs*6
PMS1
HATPase_c
mutl
MutL_Trans_hPMS_1_like
HMGB-UBF_HMG-box
932aa
TMB-low & TMB-extreme n = 33
c
TMB-high n = 30
A230Lfs*16
MSH2
MutS_I
MutS_II
MutS
MutS_IV
ABC_MSH2_euk
934aa
A230Lfs*16
R406Q
E580*
TMB-low & TMB-extreme n = 42
d
TMB-high n = 39
F1104Lfs*11
MSH6
MSH6_like
MutS_I
MutS_II
MutS
MutS_IV
MutS_V
1360aa
R976H
K1101N
R959H
K1013N
R1076H/C
E1234*
E946*
E1322*
TMB-low & TMB-extreme n = 58
e
TMB-high n = 26
D414Rfs*44/Tfs*34
V415M
R421*
PMS2
HATPase_c
mutl
MutL_Trans_hPMS_2_like
MutL_C
862aa
TMB-low & TMB-extreme n = 30

**Supplementary Fig. 8 – Landscape of mutations in *MLH1* (a), *PMS1* (b), *MSH2* (c), *MSH6* (d) and *PMS2* (e)**

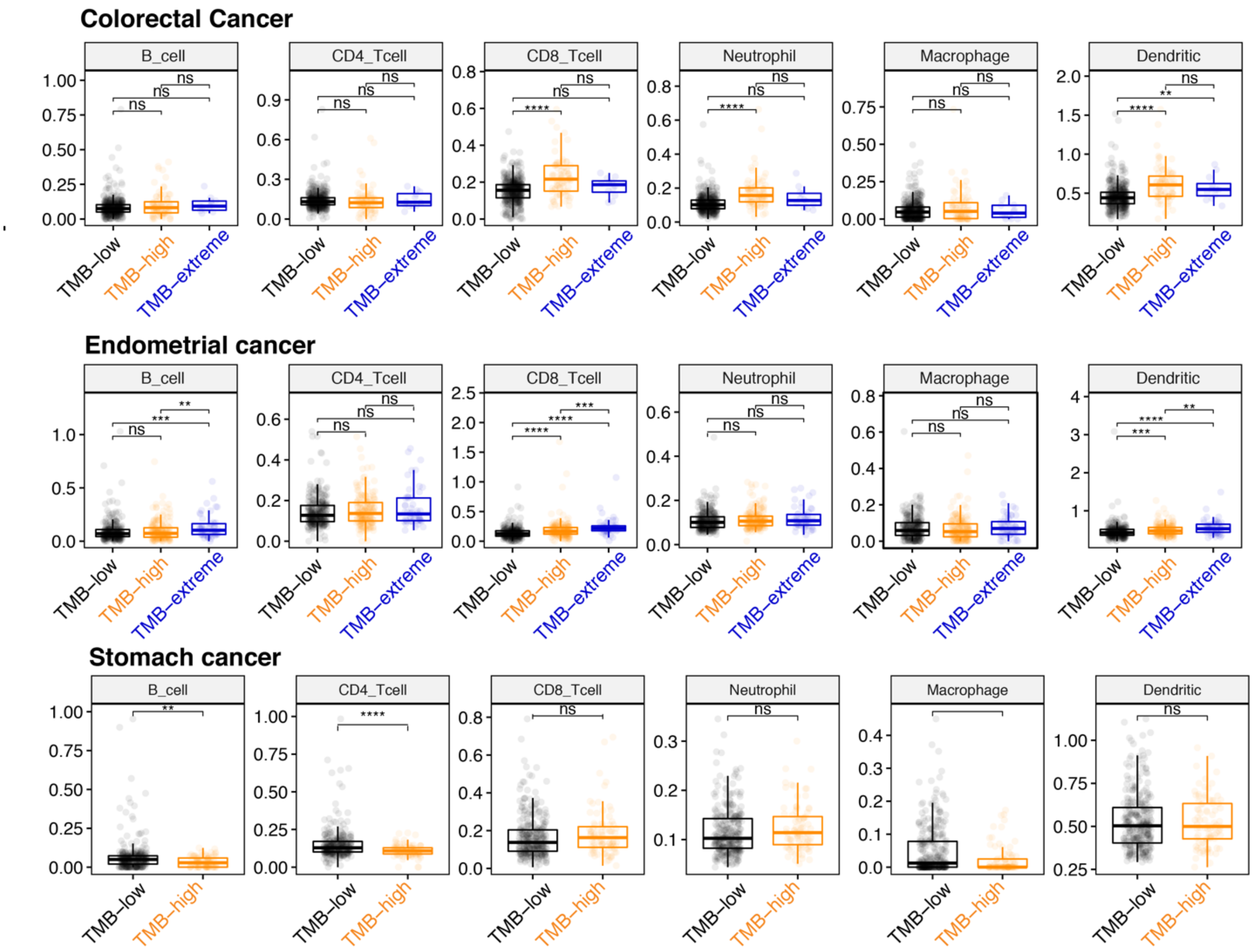


**Supplementary Fig. 9 – Abundances of immune infiltrates among three subtypes.** Box plots show the estimated abundances of tumor infiltrating immune cells among TMB subtypes in colorectal and endometrial cancer. The upper and lower quartiles of the box plots are the 75th and 25th percentiles, respectively. The whisker top and bottom are 90th and 10th percentiles, respectively. The horizontal line through the box is median value. Wilcoxon rank test was used to test the significance of the differences. The abundance of infiltrating B cell is significantly higher in only the TMB-extreme subtype compared to TMB-high and TMB-low. All differences are significant by two-sided Wilcoxon rank test in endometrial cancer but not significant in TMB-extreme subtype of colorectal cancer, which may be due to the small sample size (n = 12).

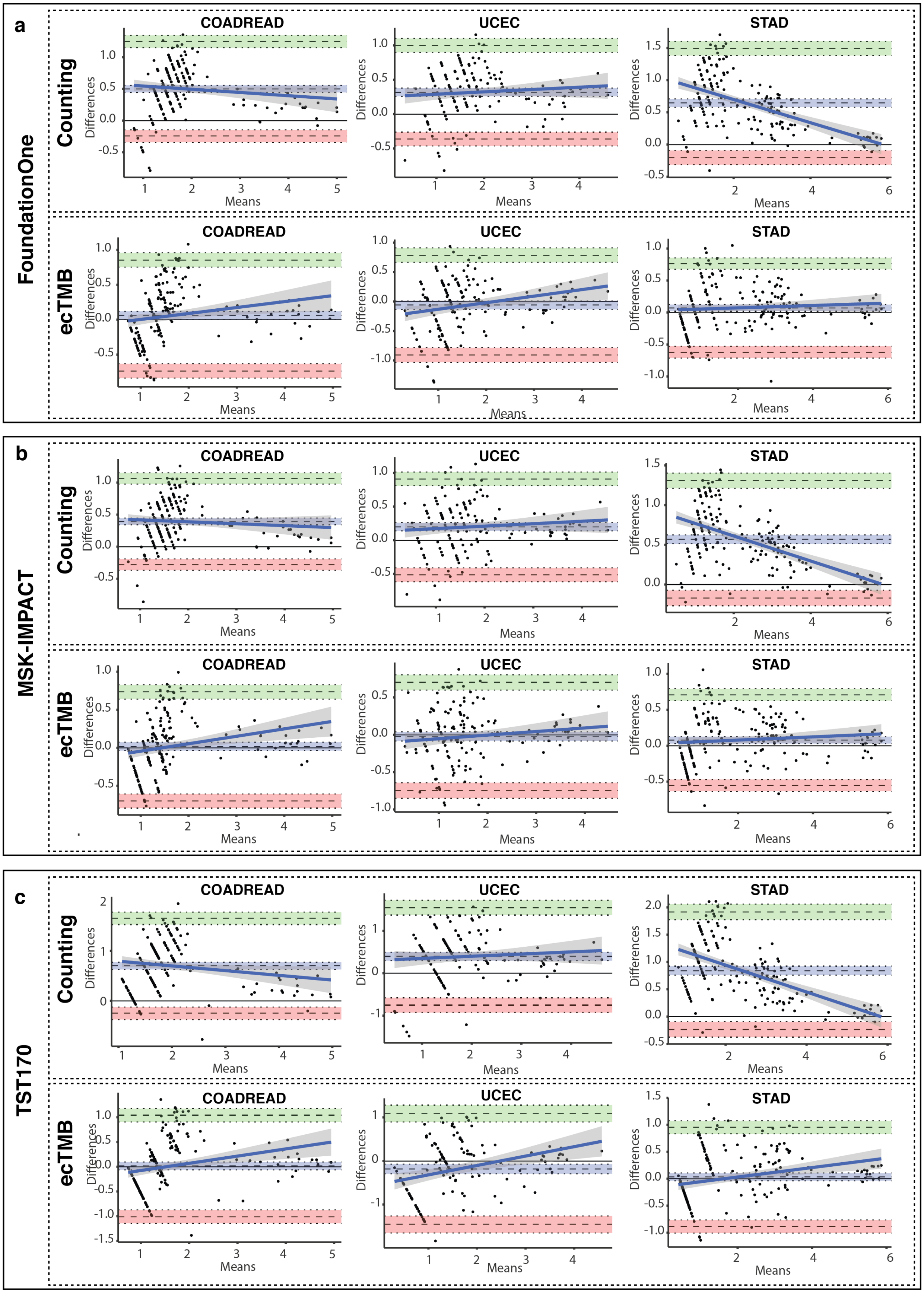

a
FoundationOne
Counting
ecTMB
b
MSK-IMPACT
Counting
ecTMB
c
TST170
Counting
ecTMB
COADREAD
UCEC
STAD
Differences
Means

**Supplementary Fig. 10 – Bland Altman plots for predictions.** (a-c) Mean of predicted panel-based TMB and standard WES-based TMB for each sample is plotted against its difference. The dashed line in the center of purple area indicates the bias (mean difference) and the purple area indicates the 95% confidence interval of bias. The green area shows the upper limits and its 95% confidence interval and the red area shows the lower limits and its 95% confidence interval. The Bland Altman analyses were done for FoundationOne (a), MSK-IMPACT (b), and TST170 panels (c). The predictions made by counting method are shown on top and ecTMB on bottom.

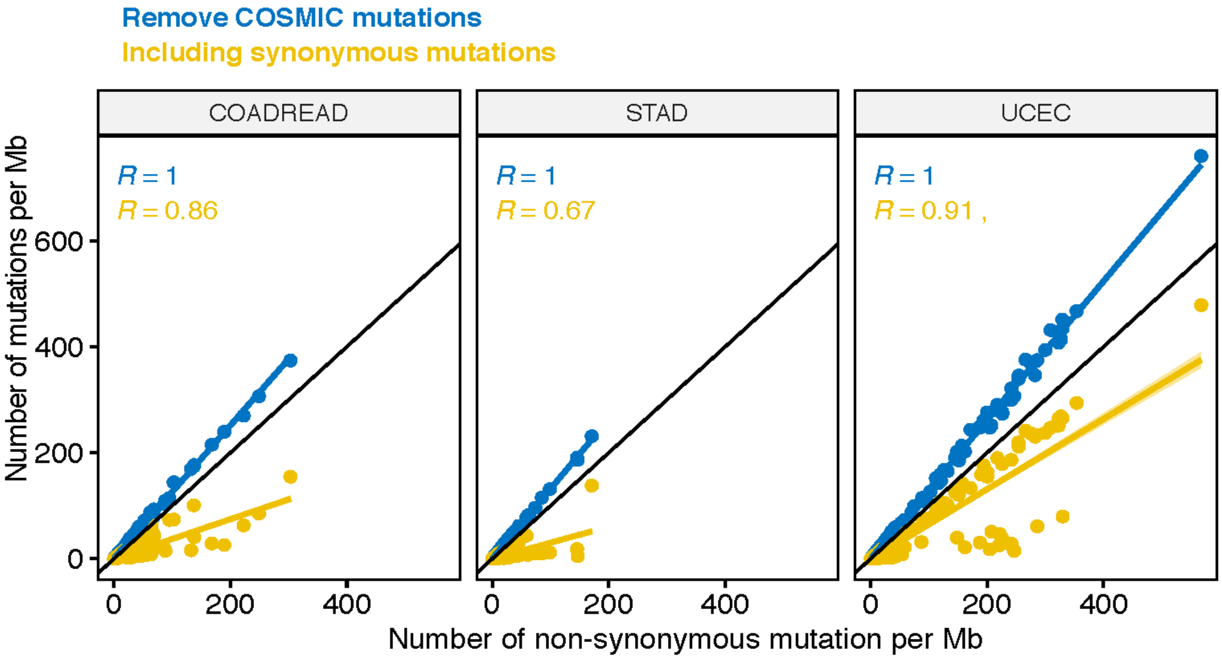


**Supplementary Fig. 11 –Correlation of WES-based standard TMB with TMB by various counting rules.** The scatter plots compare WES-based standard TMB with TMB predicted by counting non-synonymous mutations after removing COSMIC variants (blue) or adding synonymous mutations (yellow).
